# Proximity proteomics reveals a co-evolved LRRK2-regulatory network linked to centrosomes

**DOI:** 10.1101/2024.09.30.613622

**Authors:** Marita Eckert, Pasquale Miglionico, Francesca Izzi, Natalia De Oliveira Rosa, Benjamin Riebenbauer, Marius Ueffing, Francesco Raimondi, Christian Johannes Gloeckner

**Affiliations:** German Center for Neurodegenerative Diseases, 72076 Tübingen, Germany; Laboratorio di Biologia Bio@SNS Scuola Normale Superiore, 56126 Pisa, Italy; Center for Ophthalmology, Institute for Ophthalmic Research, University of Tübingen, 72076 Tübingen, Germany; Graduate School of Cellular and Molecular Neuroscience, University of Tübingen, 72074 Tübingen, Germany

**Keywords:** LRRK2 interactome, proximity proteomics, BioID, Parkinson’s disease, co-evolutionary analysis

## Abstract

Leucine-rich repeat kinase 2 (LRRK2) not only plays a vital role in familial forms of Parkinson’s disease (PD) but also represents a risk factor for idiopathic PD. Its multi-domain architecture enables fine-tuned regulation of its biological function by orchestrating intra- and intermolecular interactions. Here, we present BioID proximity proteomes of LRRK2 that reveal new interactors, which we further characterize using a novel evolutionary and structural bioinformatics pipeline. Co-evolutionary analysis of the protein-protein interaction network identifies a structural and functional module enriched in cytoskeletal components associated with the centrosome and microtubules. In addition, structural modelling of binary interactions using AlphaFold-Multimer reveals distinct groups of interactors that engage LRRK2 in a manner dependent on specific conformations and epitopes. Furthermore, we identify distinct changes in the LRRK2 proximity proteome that are induced by the type I kinase inhibitor MLi-2 or by co-expression of the LRRK2 upstream effector RAB29. Depending on its activity state and conformation, these protein-protein interactions link LRRK2 to defined cellular sub-compartments, including centriolar satellites and vesicular sub-compartments.

**Summary:** This study defines a comprehensive, LRRK2 interaction network by combining proximity proteomics with evolutionary and structural analyses, revealing how LRRK2 conformation and activity state dictate distinct protein-protein interactions. These findings link LRRK2 to conserved centrosomal, cytoskeletal, and vesicular pathways, providing mechanistic insight into its role in Parkinson’s disease.

**Key results:**

- **Centrosome-linked LRRK2 network:** BioID combined with co-evolutionary analysis identifies a highly conserved module enriched in centrosomal and microtubule-associated proteins, including CYLD as a top co-evolved interactor.
- **Conformation-specific interactions:** AlphaFold-Multimer modelling reveals distinct classes of LRRK2 interactors that bind different domains and conformations (“locked” vs. “unlocked”), corresponding to specific cellular functions.
- **State-dependent rewiring of the interactome:** The type I inhibitor MLi-2 selectively redirects LRRK2 to centriolar satellite proteins, whereas RAB29 co-expression shifts LRRK2 interactions toward vesicular and lysosomal pathways.

## Introduction

Parkinson’s disease (PD) is the second most common neurodegenerative disease after Alzheimer’s. It occurs mostly sporadically and is distributed with high frequency in the older generations. PD is characterized by a loss of dopaminergic neurons in the *substantia nigra* leading to the motor symptoms associated with the disease (Fahn, 2003). Several therapies targeting the symptoms, such as dopamine substitution, are used; however, no causative therapy currently exists. About 5–10% of PD cases are hereditary (Gasser *et al*, 2011). *LRRK2*, localized within the PARK8 locus and coding for the Leucine-rich repeat kinase 2, is one of the genes associated with familial forms of the disease but also represents a risk factor in idiopathic PD (Paisan-Ruiz *et al*, 2004; Simon-Sanchez *et al*, 2009; Zimprich *et al*, 2004). The multi-domain protein LRRK2 is an active protein kinase and, at least, mutations with confirmed disease segregation, including the most common G2019S variant within its kinase domain, augment its kinase activity (Kalogeropulou *et al*, 2022). This makes LRRK2 an important drug target (Kluss *et al*, 2022b). ATP-competitive LRRK2-specific kinase inhibitors for PD-treatment are under development and in clinical trials (Jennings *et al*, 2022). Nevertheless, structural changes of the lung observed in non-human primates indicate a relatively small therapeutic window and additionally challenge a long-term treatment with these compounds necessary to allow a life-long reduction of augmented LRRK2 pathway activation (Miller *et al*, 2023). For this reason, the identification of alternative drug targets within the LRRK2-associated pathways would be a potential solution for a better control of pathophysiological alterations associated with pathogenic forms of this protein, minimizing the risk of drug-induced lung pathologies. In addition, a larger group of patients beyond LRRK2 mutation carriers might benefit from targeting more common PD-associated pathways.

Various studies have demonstrated that LRRK2 functions in vesicle trafficking, including presynaptic vesicles as well as autophagy and lysosomal pathways (Piccoli *et al*, 2011; reviewed in: Follett & Farrer, 2021). In agreement with these cellular functions, LRRK2 phosphorylates a subset of Rab proteins, small G-proteins involved in vesicular transport, such as Rab8a and Rab10, which is elevated by PD-associated LRRK2 variants (Steger *et al*, 2017).

In common with other Roco proteins, LRRK2 features a remarkable complexity (Wauters *et al*, 2019). In fact, LRRK2 contains multiple scaffolding domains, not only involved in maintaining LRRK2 in an auto-inhibited state, as recently demonstrated by high-resolution structures of LRRK2 (Deniston *et al*, 2020; Myasnikov *et al*, 2021), but also in mediating protein–protein interactions, making LRRK2 a signaling hub possibly integrating multiple up- and down-stream signals (reviewed in: Manzoni *et al*, 2015). These studies suggest that the Roco protein LRRK2 is a tightly controlled protein kinase serving as hub and signal integrator and is most likely involved in context-specific, spatially controlled cellular responses that appear evolutionarily conserved. Considerable effort has therefore been spent on the investigation of the protein-protein interaction networks of LRRK2, mainly using affinity-based methods. Various studies have investigated the LRRK2 interactome across different cellular contexts and model organisms (Islam et al, 2016; Meixner et al, 2011; Piccoli et al, 2011; Piccoli et al, 2014). These data have been curated in public databases such as IntAct or STRING and subjected to meta-analysis to dissect functional LRRK2 signaling networks (Gloeckner & Porras, 2020; Manzoni *et al*, 2015; Zhao *et al*, 2024; Zhao *et al*, 2023). However, most approaches rely on immunoprecipitation or pull-down methods, which are biased toward stable complexes and may underrepresent transient interactions.

To overcome these limitations, we applied the BioID proximity labeling approach to identify LRRK2 interaction partners in living cells. Using BioID with three different tags, we define the proximity proteome of LRRK2 under basal conditions. In addition, using a newly designed bioinformatics workflow that estimates co-evolution of interacting partners based on their co-occurrence across sequenced genomes, we show that the resulting BioID datasets are enriched for interactions with high co-evolution scores, comparable to confirmed direct interactions. This approach identifies a cluster of interactors with strong co-evolution to LRRK2, enriched in cytoskeletal proteins associated with centrosomal and ciliary dynamics. Furthermore, structural modeling of binary complexes using AlphaFold-Multimer reveals distinct conformations and interaction interfaces linked to specific functional processes. Finally, to further stratify the LRRK2 interactome, we analyzed experimentally defined conformational states, including co-expression with the upstream effector RAB29 and stabilization of an active-like kinase conformation by a type I inhibitor (Taylor et al, 2020; Wauters et al, 2019). Notably, MLi-2, but not GZD-824, enriched a specific sub-interactome associated with centriolar satellites.

## Results

### Definition of cellular LRRK2 proximity proteomes with BioID1, BioID2 and miniTurbo

To characterize the LRRK2 interactome, we applied the proximity labeling approach BioID, which enables the identification of protein–protein interactions in living cells, including transient interactions. To define the LRRK2 proximity proteome, we generated N-terminal fusion constructs using the original BioID (BioID1), BioID2, and miniTurbo tags.

All constructs were functionally validated to confirm biotinylation activity and to ensure that the tags did not interfere with LRRK2 function. Western blot analysis demonstrated efficient biotinylation for all constructs (Appendix Figure S1), with BioID2 and miniTurbo showing higher labeling efficiency compared to BioID1. We further confirmed that the fusion proteins retained kinase activity by assessing phosphorylation of the physiological LRRK2 substrate Rab10 at Thr73 (Appendix Figure S1). Following validation, all three constructs were used to determine their interactomes upon transient expression in HEK293T cells. To increase stringency, a tag-only control was used as reference. For relative comparisons, expression levels were adjusted in preliminary experiments and consistently verified by Western blot analysis of cell lysates. The use of a tag-only control enabled efficient exclusion of endogenously biotinylated proteins, whose labeling is inherently increased by biotin supplementation.

Data from all three proximity proteomes were analyzed using SAINTexpress and, independently, by label-free quantification (LFQ) combined with an FDR-controlled t-test implemented in Perseus (Tyanova et al, 2016). To further refine the dataset, we applied a bioinformatics pipeline incorporating co-evolution analysis of interacting proteins and structural modeling using AlphaFold-Multimer (Figure 1A).

**Figure 1:**
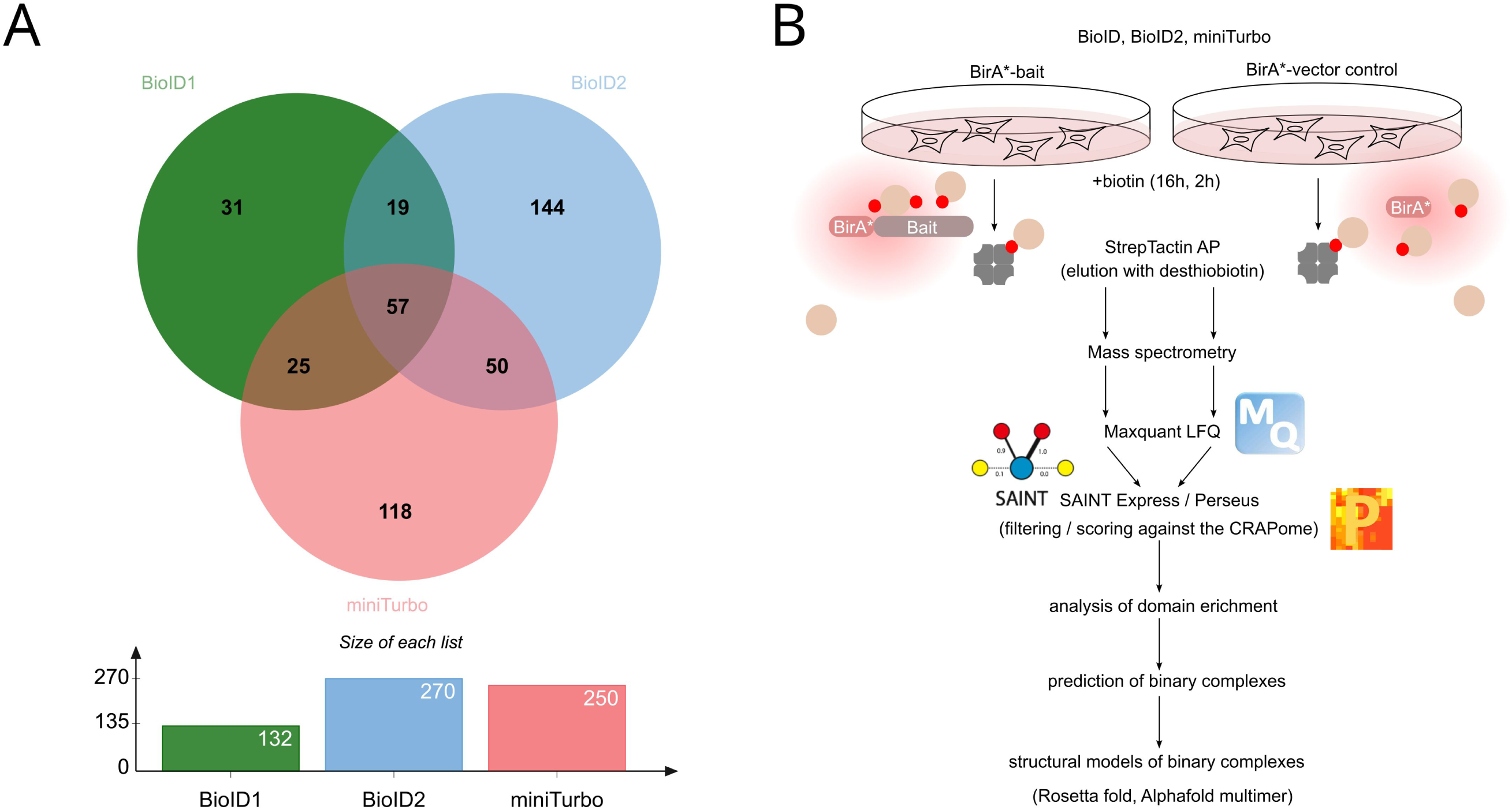
(A) Workflow of the BioID analysis and post processing of the data. (B) Overlap between the different BioID approaches. Enriched proteins were determined by relative label-free quantification (combined with an FDR-controlled T-test, P = 0.05) of BioID LRRK2 vs the corresponding tag-only control: BioID1-LRRK2/BioID1-tag: N = 8, BioID2-LRRK2/BioID2: N = 9, miniTurbo-LRRK2/miniTurbo tag: N = 8.

Using LFQ, 168, 312, and 241 proteins were identified as significantly enriched in the LRRK2 condition compared to the tag-only control for BioID1, BioID2, and miniTurbo, respectively (Figure 1B; Dataset EV1). SAINTexpress (Teo et al, 2014), followed by filtering against the CRAPome database (Mellacheruvu et al, 2013), provided a more stringent set compared to LFQ-based analysis. Based on this approach, we defined a non-redundant set of 208 unique interactors, hereafter referred to as the “BioID set,” which was subjected to downstream bioinformatics analysis.

### Functional-enrichment analysis of the LRRK2 proximity proteome

We compared the LRRK2 BioID interactome with a reference LRRK2 interactome available from IntAct (Orchard *et al*, 2014). We found that only 54 interactors in the filtered BioID set were previously reported to be associated with LRRK2, whereas the majority (154) have not been reported before (Figure 2A, B). By integrating the LRRK2 BioID interactome with known interactions from IntAct, we defined modules of proteins that interact not only with LRRK2 but also with each other, thereby revealing structured interaction networks (Figure 2A). Among the enriched intracellular and cytosolic compartments, we found terms related to microtubules (e.g. Microtubule Cytoskeleton, FDR = 5.83E-15) and Centrosome (FDR = 1.43E-13), emphasizing the importance of LRRK2 in microtubule organization (Figure 2C), and representing the largest module of interacting proteins in the network (Figure 2A). In addition, we also found processes related to cell cycle and mitosis (e.g. “Regulation of mitotic cell cycle”, FDR = 9.71E-08) (Figure 2D).

**Figure 2:**
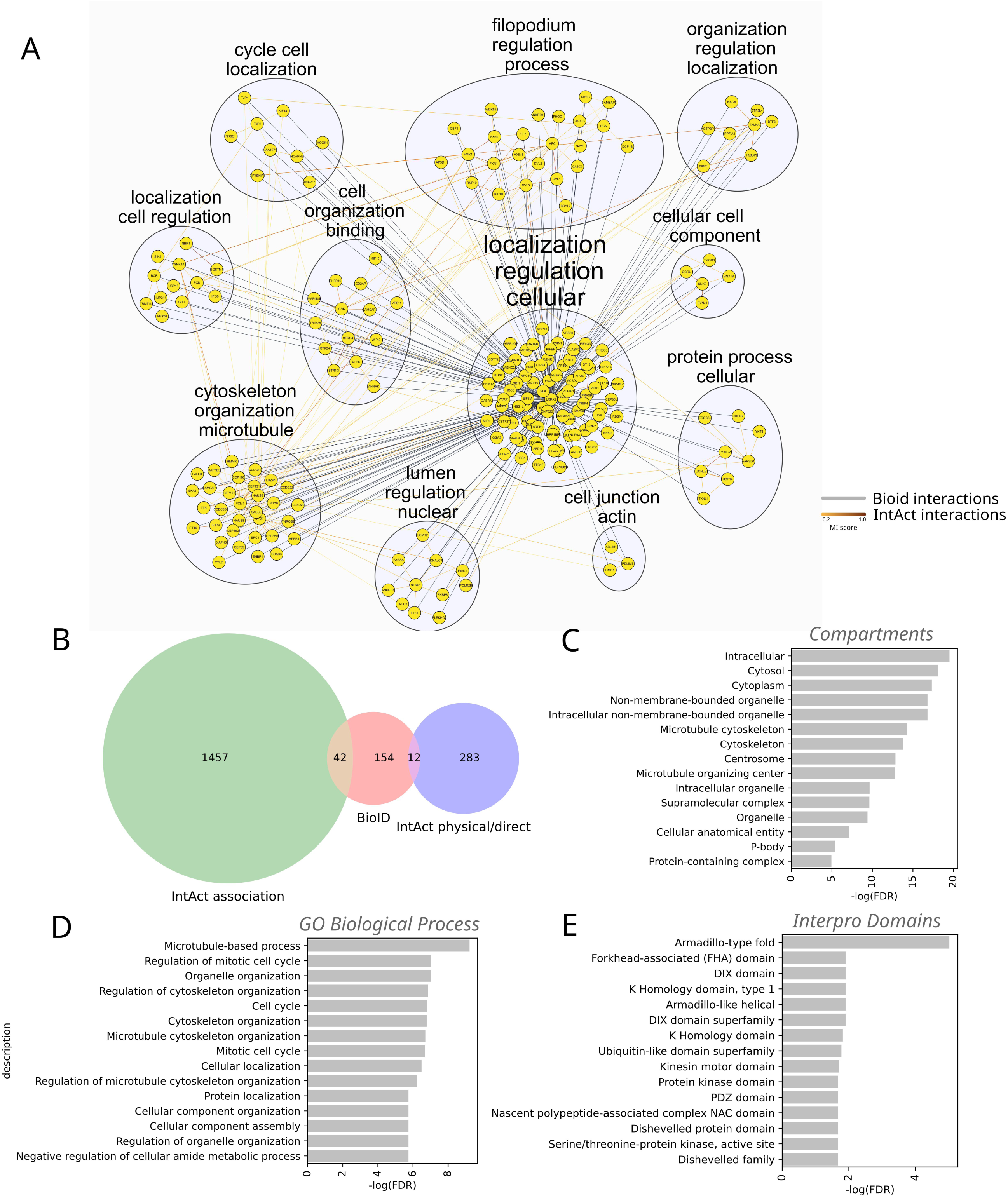
LRRK2 interactome functional enrichment. A) the LRRK2 BioID interaction network was mapped on known PPIs from IntAct via Cytoscape. The edges’ color and thickness for mapped interactions are proportional to IntAct MI score, else they are colored in gray with constant thickness. Modules within the network have been identified through the Glay method of the Autoannotate app and nodes colored accordingly. Labels were calculated using the ‘Adjacent Words’ setting of the WordCloud algorithm, based on significantly enriched (FDR < 0.01) Gene Ontology (GO) Biological Processes and Cellular Component terms associated with each gene in the network; B) Venn diagram showing the overlap between the BioID interaction sets and known LRRK2 interactions in IntAct; barplots showing the top 15 most significant terms obtained through String’s functional enrichment of C) Compartments, D) GO Biological Process and D) Interpro domains.

Multiple protein domains instances are significantly enriched within the list of identified LRRK2 interaction partners (Figure 2E). In particular, we found that the Armadillo-type fold instance is the most enriched one (FDR = 9.58E-06), which is found in 21 distinct interactors, followed by other domains with lower, but still significant, degree of enrichment, such as “Forkhead-associated (FHA) domain” (Almawi *et al*, 2017), found in six distinct proteins, the “DIX domain” (Ehebauer & Arias, 2009; Kafka *et al*, 2014) found in four proteins, the “K Homology domain, type 1” (Grishin, 2001) found in six proteins and “Ubiquitin-like domain superfamily” (FDR < 0.05), the latter being found in 11 distinct interacting proteins. Interestingly, LRRK2 and its Drosophila orthologue dLRRK have been demonstrated to functionally interact with the Forkhead box transcription factor FoxO1/dFoxO. In particular, LRRK2-mediated phosphorylation of FoxO1 enhances its transcriptional activity leading to neuronal toxicity (Kanao *et al*, 2010).

### A cluster of proteins involved in centrosome/ microtubular dynamics are most co-evolved with LRRK2

We analyzed the extent of co-evolution of LRRK2 with its interactors to prioritize the most conserved relationships associated to LRRK2’s functions. We derived a metric for the co-evolution between two interacting proteins based on the degree of co-presence, assessed through a Jaccard score, of interacting gene pairs in sequenced genomes available from a reference orthology database (e.g. OMA, see Methods). We found that such simple co-evolutionary metric is informative about non-enzymatic, direct interactions as well as stable complexes at a human interactome-wide level (Meldal *et al*, 2022; Miglionico *et al*, 2024; Orchard *et al*, 2014) (Dataset EV2A). We found similar trends when focusing on LRRK2 interactions reported in IntAct, with “non-enzymatic direct” PPIs being characterized by a significantly higher Jaccard score compared to those categorized as “association” or “physical interaction” (Figure 3A). Intriguingly, we found that the distribution of co-evolution metrics calculated for BioID interactors is significantly higher than most IntAct categories (i.e. “association”, “physical”, “enzymatic direct”) and it is as high as the non-enzymatic direct interactions (Figure 3A). This result suggests that interactors captured within the BioID dataset may represent stable, direct interaction partners of LRRK2 with whom they co-evolved. Likewise, simple co-evolution scores are predictive for LRRK2 direct interactions, particularly the non-enzymatic direct ones (Figure 3B; ROC AUC = 0.715). In the case of the LRRK2 interactome, we found that this co-evolutionary metric is more predictive of non-enzymatic direct interaction compared to another co-evolution estimate, i.e. Mutual Information (ROC AUC = 0.66; Appendix Figure S2), which is instead more predictive at a human proteome scale (Miglionico *et al*, 2024).

**Figure 3:**
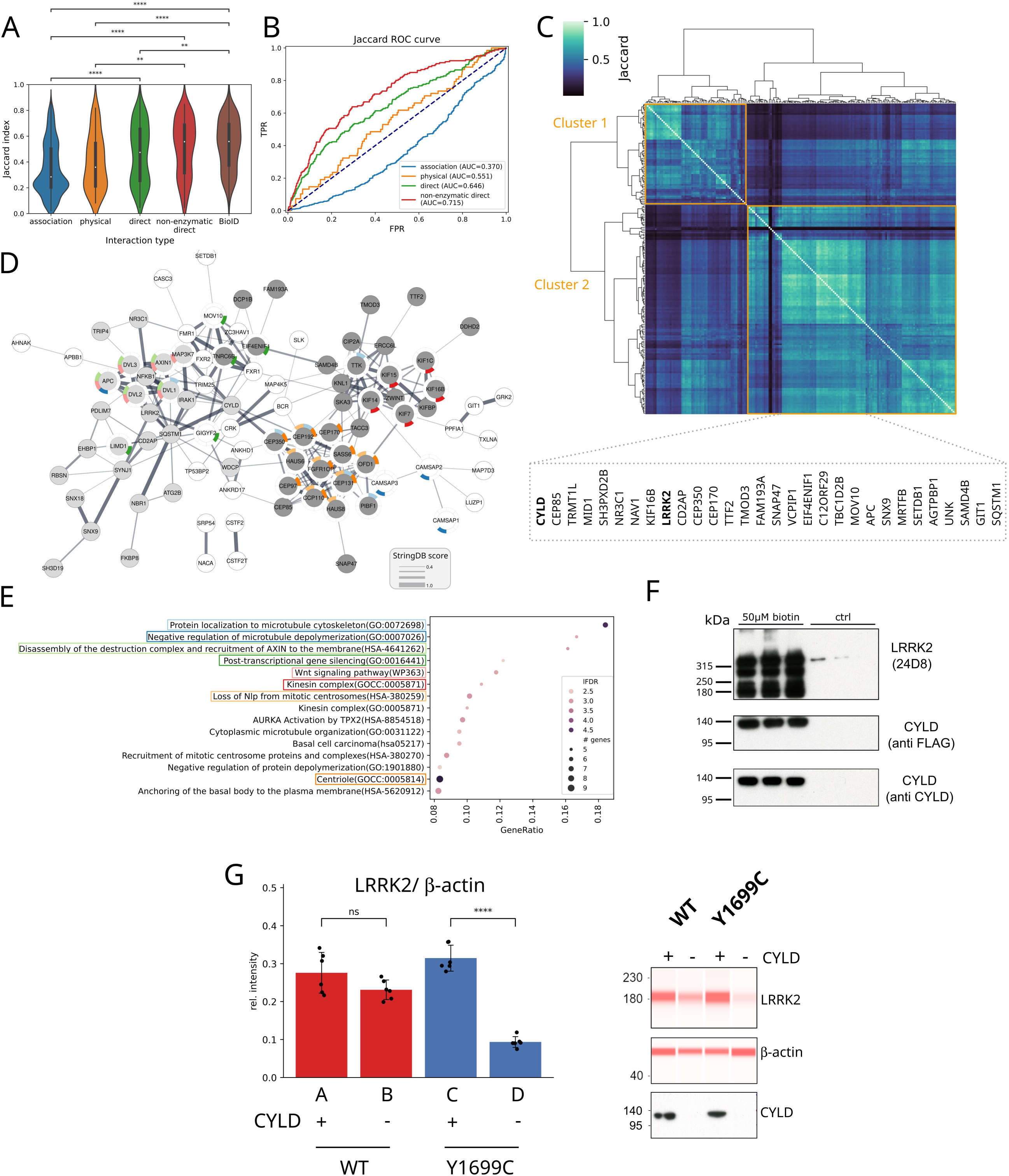
Co-evolution analysis of LRRK2 interactome. (A) Distribution of co-evolution metric (i.e. Jaccard score) among the different LRRK2 interaction sets, including IntAct interaction types and BIOID. P-values have been computed with a two-sided Mann-Whitney test with Bonferroni correction (* *P* < 0.05, ** *P* < 0.01, *** *P* < 0.001, **** *P* < 0.0001). The plot shows the comparison between 2062 independent samples divided in 5 classes with the following number of elements: association 1424, physical 68, direct 215, non-enzymatic direct 166, BioID 189. P-values: association-direct: 1.2E-11; association-non-enzymatic direct: 5.8E-19; association-BioID: 6.1E-29; physical-non-enzymatic direct: 0.0017; physical-BioID: 2.7E-5; direct-BioID: 0.0082. Violin plots show the median as the white central dot and first and third quartiles as bounds of the thicker black line (box); the whiskers extend to the last data point within 1.5 times the interquartile range (IQR) from the box’s boundaries. (B) predictiveness of LRRK2 interaction type via co-evolution (C) Heatmap showing the hierarchical clustering based on the degree of co-evolution (Jaccard score) of the LRRK2 proximity proteome. (D) String network of the proteins most co-evolved with LRRK2 participating to cluster 2. Nodes are colored in gray or light-gray according to modules membership determined with the MCL approach available from the Cytoscape clusterMaker app. Node rims are colored according to enriched processes. (E) Dot plot showing top 15 significantly enriched (FDR < 0.01) categories from the string network of cluster 2. Values on the x axis display the ratio of genes of the term participating to the network over the total number of genes in that category. The ratio of the dot is proportional to the number of genes of that category in the network, while the node color is the darker the more significant is the category enrichment. Only categories with at least 5 genes have been displayed. Representative, enriched categories have been colored with the same color scheme as in panel (D). (F) Validation of CYLD as part of the LRRK2 proximity proteome by Western blot. Because suitable antibodies for detecting CYLD at endogenous levels were not available, a FLAG-HA–tagged CYLD construct was recombinantly expressed for this experiment. This contrasts with the MS–based approach, which relied on endogenous protein levels. Prior to antibody incubation, the membrane was horizontally cut above the 140 kDa marker (expected molecular weight of CYLD: ∼110 kDa). The upper part of the blot was probed with an anti-LRRK2 antibody (24D8), while the lower part was probed with anti-FLAG and subsequently re-probed with anti-CYLD antibodies. (G) Effect of CYLD co-expression on steady-state LRRK2 levels: LRRK2 WT or Y1699C was co-expressed with CYLD or empty vector control. LRRK2 protein levels were normalized by β-actin. Statistical significance was assessed using unpaired two-tailed t-tests (P-values: A-B: 0.0951; C-D: 4.5E-8; N = 6 biological replicates; error bars = SD) (left panel). Reconstructed Capillary Western signals and confirmation of CYLD expression by Western blot. On the capillary Western blot, LRRK2 migrates at a lower molecular weight (right panel).

We used the co-evolution metric to cluster LRRK2 and its interactors, which revealed groups of proteins characterized by reciprocal high co-evolution (Figure 3C). The different groups of interactors are also characterized by specific biological processes, which are overall mutually exclusive to each other, suggesting that clusters of co-evolved proteins underpin specific functional interactions and biological processes (Appendix Figure S3). Notably, the cluster containing LRRK2 (i.e. Cluster 2) is characterized by on average higher co-evolution and several highly co-evolved proteins (Figure 3C). The top 10 proteins of Cluster 2 showing highest co-evolution are summarized in Table 1. By mapping members of this cluster onto the STRING functional interaction network (Figure 3D, see Methods), we found functional interactions for most of the members, including LRRK2. Overall, STRING yielded a highly significant PPI enrichment (P = 1.0E-16), suggesting that these proteins are more likely to interact with themselves than with randomly selected proteins. Moreover, network topology analysis delineated two distinct submodules: the first encompassing LRRK2 together with proteins predominantly linked to Wnt signaling (FDR = 0.0013; Figure 3E), and the second consisting of proteins implicated in cytoskeletal processes (“Protein localization to microtubule cytoskeleton”; FDR = 2.18E-05), centrosome dynamics (“Loss of Nlp from mitotic centrosomes”; FDR = 0.00086) and centriole organization (“Centriole”; FDR = 1.11E-05). Consistently, Cluster 2 displayed strong enrichment for cellular component terms, most notably “Microtubule Cytoskeleton” (FDR = 2.10E-12) and “Centrosome” (FDR = 1.01E-12) (Dataset EV2B). Interestingly, CYLD is the most co-evolved protein of LRRK2, a deubiquitinase that acts as a negative regulator of dopamine neuron survival in Parkinson’s disease (Pirooznia et al., 2022), but has not yet been reported to directly interact with LRRK2. Notably, CYLD emerges as a critical bridging element that connects the two principal submodules within Cluster 2’s interactome (Figure 3D).

**Table 1:** 10 IDs with the highest Jaccard score underlying identified “Cluster 2”. The DUB CYLD (bold) shows highest co-evolution to LRRK2.

| Accession | Gene name | Protein name | Jaccard score |
| --- | --- | --- | --- |
| Q5S007 | LRRK2 | Leucine-rich repeat serine/threonine-protein kinase 2 (bait) | 1.000000 |
| **Q9NQC7** | **CYLD** | **Ubiquitin carboxyl-terminal hydrolase CYLD** | **0.848485** |
| Q6P2H3 | CEP85 | Centrosomal protein of 85 kDa | 0.846626 |
| Q96JH7 | VCPIP1 | Deubiquitinating protein VCPIP1 | 0.834286 |
| Q96L93 | KIF16B | Kinesin-like protein KIF16B | 0.832335 |
| P78312 | FAM193A | Protein FAM193A | 0.817647 |
| Q8NEY1 | NAV1 | Neuron navigator 1 | 0.817073 |
| Q7Z2T5 | TRMT1L | TRMT1-like protein | 0.815951 |
| Q5SZL2 | CEP85L | Centrosomal protein of 85 kDa-like | 0.810127 |
| O95684 | CEP43 | Centrosomal protein 43 | 0.809816 |
| Q96RS0 | TGS1 | Trimethylguanosine synthase | 0.798742 |

On the other hand, Cluster 1 is characterized by distinct processes, such as FAR/SIN/STRIPAK complex (P = 0.0012; Dataset EV2 B). Taken together, these results suggest that our co-evolution-based approach was able to identify a group of LRRK2 interacting proteins, mostly associated to microtubule and centrosome functions via the formation of stable, supramolecular complexes.

Likewise, co-evolution-based clustering of LRRK2 interactors from IntAct shows interactors such as RAB29, RAB12, LRRK1 or SNCA among the most co-evolved partners that cluster with LRRK2 (Appendix Figure S4), confirming the tight evolutionary constraint on LRRK2’s functional regulation.

After validating CYLD as part of the LRRK2 proximity proteome by Western blot analysis (Figure 3F), we investigated the functional relevance of this association by assessing whether CYLD influences LRRK2 abundance. To this end, LRRK2 wild-type or the pathogenic Y1699C variant was co-expressed with CYLD or an empty vector control. Y1699C was chosen, as this variant shows a robust increase in Rab phosphorylation (Kalogeropulou *et al*, 2022) and reduced steady-state protein levels under our experimental conditions. As classical ECL-based Western blot did not allow reliable quantification of the large protein LRRK2 (286 kDa), we used a capillary-based Western system with fluorescent detection.

CYLD co-expression was associated with increased steady-state LRRK2 levels across two independent experiments (N = 6 biological replicates each), with the effect being more pronounced for the Y1699C variant. Wild-type LRRK2 showed a smaller increase, consistent with a weaker trend (Figure 3G; Appendix Figure S5A).

Because CYLD preferentially acts on K63-linked chains, we repeated the experiment with additional co-expression of either wild-type ubiquitin or a K63-only ubiquitin mutant, which restricts linkage to lysine 63 (Lim *et al*, 2005). However, while replicating the effects seen without Ubiquitin overexpression, these data do not allow conclusions about the contribution of specific ubiquitin linkage types (Appendix Figure S5B).

### AF-based modelling reveals a discrete repertoire of conformations and interfaces shaping the LRRK2 interactome

We predicted 3D complexes for binary interactions of LRRK2 and 196 partners in the BioID dataset via AlphaFold-multimer (AF) and characterized their interfaces and conformations (Figure 4A).

**Figure 4:**
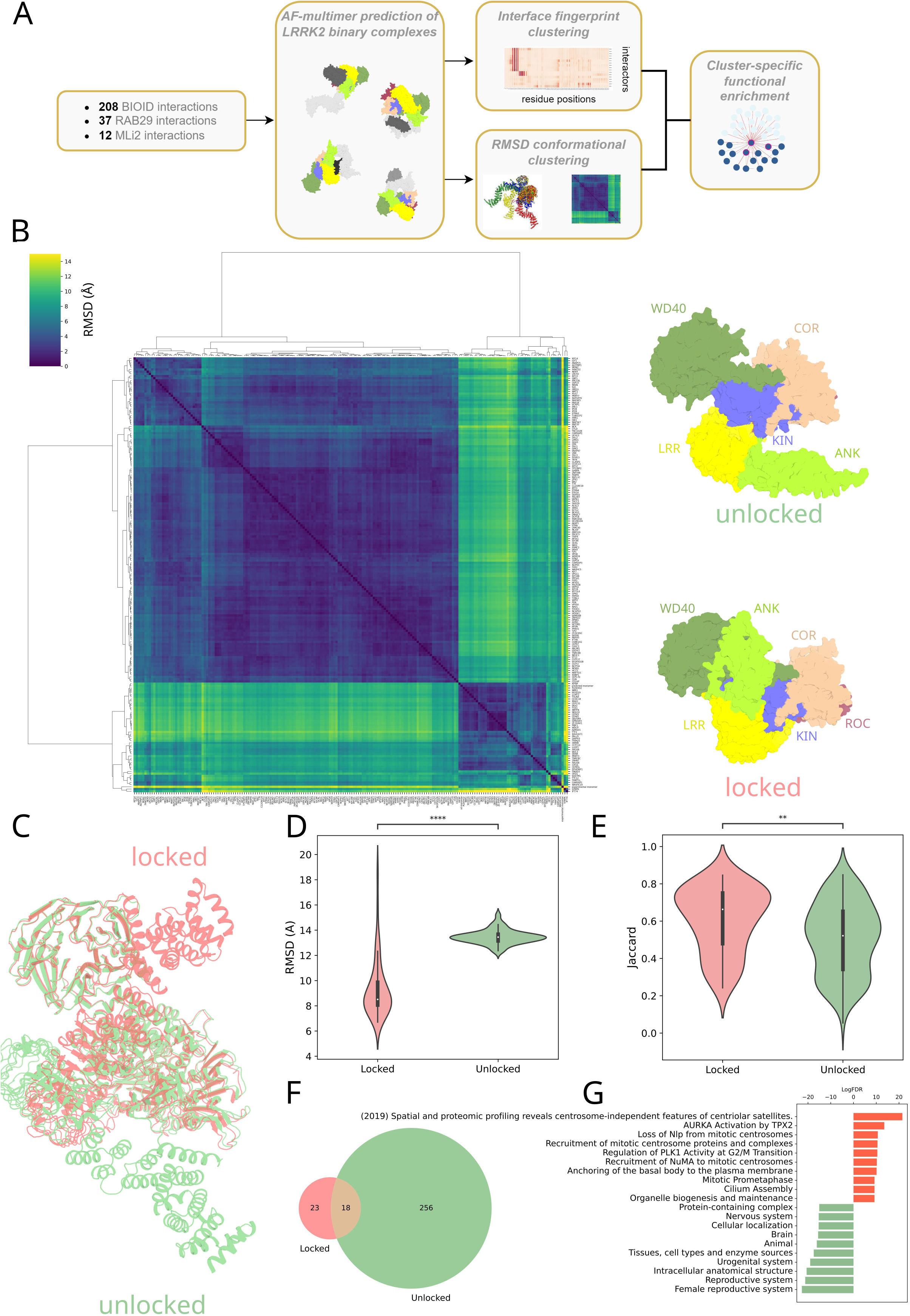
3D complexes prediction and analysis of LRRK2 interactome (A) workflow of the structural bioinformatics analysis of LRRK2 interactome; (B) on the left the heatmap showing the hierarchical clustering based on pairwise RMSD calculation of LRRK2’s chain in the different complexes. On the right two representative conformations of LRRK2 in the “locked” and “unlocked” conformation to mark the two main clusters; (C) cartoon representation of “locked” (tomato) and “unlocked” (light green) superimposed LRRK2 conformations; (D) violin plot of the distribution of RMSD calculated with respect to the reference LRRK2 experimental structure (PDB 7LHW) for the locked and unlocked clusters (P = 3.8E-20; comparison between 48 locked and 148 unlocked independent samples). Color scheme as in (C); (E) distribution of the Jaccard score for the locked and unlocked clusters (P = 0.0013; comparison between 48 locked and 148 unlocked independent samples). Color scheme as in (C). P-values for violin plots in D and E have been computed with a two-sided Mann-Whitney test (* *P* < 0.05, ** *P* < 0.01, *** *P* < 0.001, **** *P* < 0.0001). Violin plots show the median as the white central dot and first and third quartiles as bounds of the thicker black line (box); the whiskers extend to the last data point within 1.5 times the interquartile range (IQR) from the box’s boundaries; (F) Venn diagram of the enriched terms in the locked and unlocked clusters. Color scheme as in (C); (G) Top 10 exclusively enriched (FDR < 0.01) terms in the locked and unlocked clusters.

We structurally aligned LRRK2 conformations in the different complexes by superimposing the monomeric LRRK2 chains and then calculating the pairwise Root Mean Squared Deviation (RMSD) of all the possible pairs (Figure 4A; Methods). We performed hierarchical clustering of LRRK2 structures based on RMSD, through which we identified two main clusters (Figure 4B). A smaller one is characterized by conformations with lower deviation from the inactive LRRK2 structure (PDB ID: 7LHW), which we termed “locked” (Figure 4B-D). The main structural hallmarks characterizing the “locked” conformation, which are inherited by the experimental inactive LRRK2 structure (PDB:7LHW) (Myasnikov *et al*, 2021), are the LRR wrapping around the catalytic domains, particularly the kinase, and a contact formed by the extended loop within the LRR repeats, which includes the hinge helix, and the C-terminal WD40 (Myasnikov *et al*, 2021). A larger cluster comprised conformations characterized by a greater deviation from the inactive structure due to major rearrangements of the LRR, ROC and COR domains, which we termed “unlocked” (Figure 4B-D). Such conformations are reminiscent of the ones that we observed in the predicted LRRK2-RAB10 complex, which we validated as the AF-predicted conformation best accommodating experimental XL-MS data (preprint: Guaitoli *et al*, 2023). Intriguingly, we found that proteins interacting with LRRK2 in these two alternative conformations are associated with distinct evolutionary and functional properties. Indeed, LRRK2’s complexes in the “locked” conformations are characterized by a significantly lower RMSD (P = 3.81E-20; Figure 4D) and higher co-evolution (P = 1.33E-3; Figure 4E) than the complexes in the “unlocked” conformations. The two groups are also characterized by largely exclusively enriched processes (Figure 4F). In particular, the interactors engaging with LRRK2 in the locked conformation are significantly associated with processes linked to centriolar satellites (P = 4.76E-10) and cilium assembly (P = 0.0001) (Figure 4G; Dataset EV2C). Those interacting with the unlocked conformations are associated with a higher number of enriched processes, including cluster-specific ones such as cytoskeleton (P = 4.53E-09) or the microtubule organizing center MTOC (P = 6.40E-07). Hence, the predicted LRRK2 conformational switch might be relevant in orchestrating the interaction with partners associated with distinct functions.

We also explored patterns of engagement of interactors with the different regions of the LRRK2 multi-domain architecture. To this end, we considered the distribution of distance probabilities (i.e. distograms) generated by AF for each binary interaction and mapped the most likely interacting residues of the partners with respect to LRRK2 positions. We used these partner-specific interaction fingerprints to cluster the interactome and identified four major groups of proteins, each named according to the LRRK2 3D interface involved in the interaction (Figure 5A; Dataset EV2 D; see Methods). The Core cluster comprises partners that interact with the LRR–ROC–COR–KIN domains and was named accordingly, as these domains represent the most evolutionarily conserved regions of LRRK2 (Figure 5A, B). The WD40 cluster includes complexes involving the homonymous domain (Figure 5A, C), while the ARM cluster comprises proteins engaging a newly identified site within the Armadillo domain (Figure 5A, D). Lastly, the “Other” cluster contains proteins with alternative interaction modalities or no clearly defined binding interface (Figure 5A). Complexes predicted to interact with the Core group displayed significantly higher AlphaFold confidence scores compared to those interacting with the WD40 domain (P = 1.14E-8) or via “Other” interaction modalities (P = 4.86E-14; Figure 5E). Complexes of the Core group show a significantly higher RMSD of the LRRK2 chain compared to the WD40 group (P = 4.05E-3; Figure 5F), whose interactors bind LRRK2 in both locked and unlocked conformations (see below). In the Core cluster we found established LRRK2 interactors, such as DVL1-2, confirming the known interaction mechanism through the ROC-COR domains (Sancho *et al*, 2009). In the same cluster, we also found novel interactors discovered within the BioID set, such as e.g. CYLD interacting through the Core interface in a similar way. A second cluster (WD40) displayed higher likelihood of interaction with the WD40 domain (Figure 5A, C), with predicted complexes characterized by a similar interface involving the binding of a short, disordered peptide stretch to a conserved crevice formed by β-propeller loops of the WD40 domain (Figure 5C). WD40 domain binders might bind to LRRK2 in either locked or unlocked conformation (Figure 5A, F), as also suggested by a bi-modal distribution of the LRRK2 chain RMSD (Figure 5F). These interactors also displayed significantly higher co-evolution with respect to “Other” LRRK2 interactors (P = 5.57E-4; Figure 5G). The ARM cluster involves proteins showing binding preferences for a new docking site on the Armadillo domain, centered around residues 500-600 and mediating the interaction with proteins such as AXIN1 (Figure 5A, D). The larger cluster (Other) contains proteins interacting with weaker likelihood and variable docking sites (Figure 5A). Among these, we also found proteins engaging with two additional ARM docking sites, including one (around aa 240) which has been recently shown to interact with RAB12 (Dhekne *et al*, 2023; Li *et al*, 2024) and mediate a RAB29-independent LRRK2 Kinase activity activation (Purlyte *et al*, 2018; Zhu *et al*, 2023), which is predicted to interact with proteins such as KIF16B. A third ARM binding region, encompassing aa 354-444, corresponds to the RAB29 interaction interface found in the recently determined RAB29-bound, tetrameric LRRK2 structure (Zhu *et al*, 2023), and it is the predicted interface for proteins such as NBR1 and MON2.

**Figure 5:**
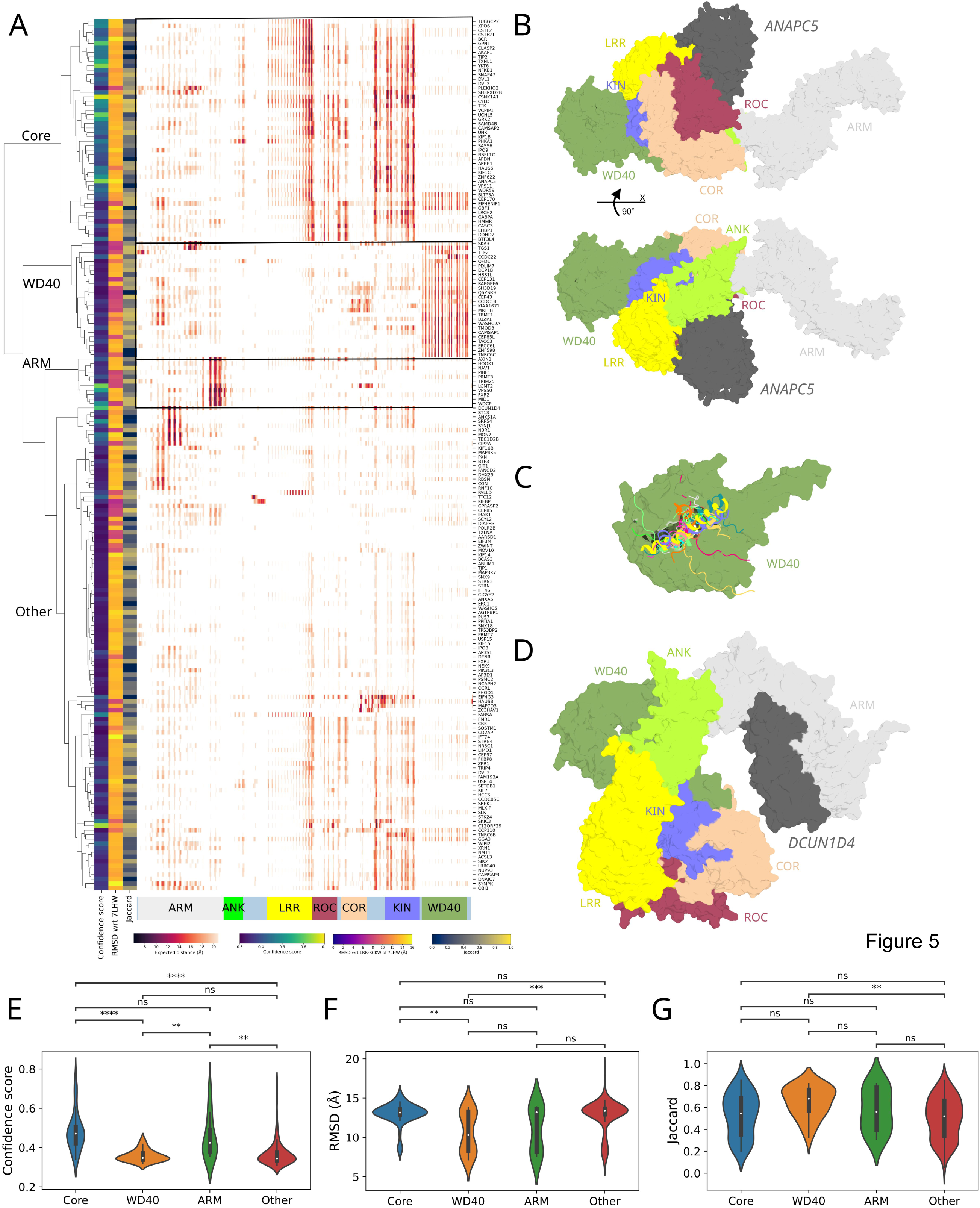
Interface analysis of LRRK2’s 3D interactome (A) interface fingerprint of the LRRK2 complexes via distogram calculation. Cells contain the minimum expected distance obtained from the distograms of any residue of each interactor (row) to every residue in LRRK2 (column). Rows are color annotated based on confidence score, Jaccard and RMSD from reference LRRK2 structure (PDB: 7LHW); (B) representative structure of the Core cluster (i.e. LRRK2-ANAPC5 complex); (C) representative structures of the WD40 cluster; (D) representative structures of the ARM binding mode cluster (i.e. LRRK2- DCUN1D4 complex); distribution among different interface cluster of the (E) confidence score, (F) RMSD and (G) Jaccard score descriptors. (C), (D) and (E) show the comparison between 196 independent samples divided in 4 cluster with the following number of elements: 50 Core, 26 WD40, 11 ARM and 109 Other. P-values have been computed with a two-sided Mann-Whitney test with Bonferroni correction (* *P* < 0.05, ** *P* < 0.01, *** *P* < 0.001, **** *P* < 0.0001). Violin plots show the median as the white central dot and first and third quartiles as bounds of the thicker black line (box); the whiskers extend to the last data point within 1.5 times the interquartile range (IQR) from the box’s boundaries. (C) P-values: Core-WD40: 1.1E-8; Core-ARM: 1.0; Core-Other: 4.9E-14; WD40-ARM: 0.0035; WD40-Other: 1.0; ARM-Other: 0.0030. (D) P-values: Core-WD40: 0.0040; Core-ARM: 1.0; Core-Other: 1.0; WD40-ARM: 1.0; WD40-Other: 5.6E-4; ARM-Other: 1.0. (E) P-values: Core-WD40: 0.099; Core-ARM: 1.0; Core-Other: 1.0; WD40-ARM: 1.0; WD40-Other: 0.0026; ARM-Other: 1.0.

Overall, these results further support the hypothesis that interactors *de-novo* identified by proximity labelling may physically interact with LRRK2 domains through specific conformations and contact interfaces. Interactors lacking a strong interaction probability in the distogram might represent proteins interacting with LRRK2 either transiently or indirectly.

### Type I but not type II inhibitors lead to conformational changes directing LRRK2 to centriolar satellites

As mentioned above, LRRK2 adopts multiple conformations through which it engages with its interactors. Effector-mediated oligomerization to an asymmetric tetramer has been demonstrated to switch LRRK2 to an active conformation (Zhu *et al*, 2023). In addition, ATP-competitive kinase inhibitors can stabilize different conformations. While type I inhibitors stabilize the kinase domain in an active-like conformation, type II inhibitors stabilize the inactive conformation of the kinase domain (Deniston *et al*, 2020; Raig *et al*, 2025; Zhu *et al*, 2024). Interestingly, type I but not type II inhibitors lead to a rapid dephosphorylation of LRRK2 phospho-sites in an interdomain space downstream of the hinge-helix which locks LRRK2 in a compact inactive fold by binding the C-terminal WD40 domain (Kalogeropulou *et al*, 2022; Myasnikov *et al*, 2021). We hypothesized that specific inhibitors could distinguish the proximity proteome of LRRK2 in its active versus inactive conformation. To test this, we compared the LRRK2 proximity proteome in HEK293 cells stably expressing miniTurbo-tagged LRRK2 following treatment with the type I kinase inhibitor MLi-2, the type II inhibitor GZD-824 (previously shown to inhibit LRRK2 in cells (Kalogeropulou *et al*, 2022; Tasegian *et al*, 2021)), or a DMSO control. Western blot analysis of LRRK2 pS935 and pT73-Rab10 confirmed effective inhibition under these conditions (Appendix Figure S6A). Quantitative label-free analysis combined with an FDR-controlled t-test revealed that only MLi-2 treatment led to the enrichment of a distinct set of proteins associated with cilia assembly and centriolar satellites in the LRRK2 proximity proteome (Figure 6A–E, Appendix Figure S6B), a pattern not observed with GZD-824. Intriguingly, interactors determined in the presence of the MLi-2 inhibitor are characterized by a significantly higher co-evolution (Figure 6F) compared to other BioID interactors. The complete list of significantly enriched proteins (MLi-2 vs. DMSO) is shown in Table 2. Additional information is provided in Dataset EV1.

**Figure 6:**
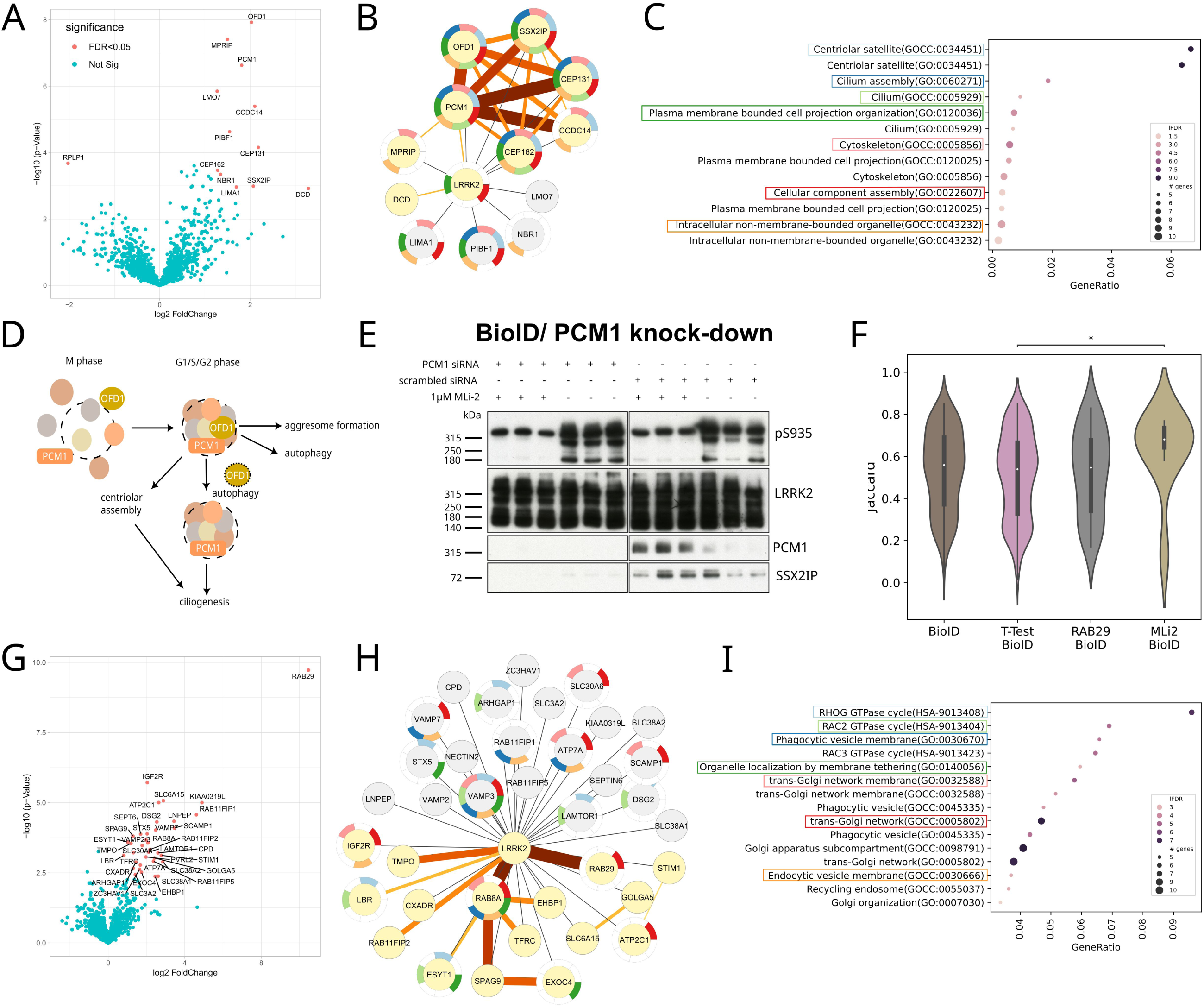
Different effects of type I and type II inhibitors and RAB29 on the LRRK2 proximity proteome. (A) Volcano plot MLi-2 vs DMSO (moderated permutation-based T-test, FDR = 0.05; S0 = 0.1, N = 7). (B) Stringified network of the MLi-2 interactome from (A) mapped to IntAct known interactions. Edges’ color and thickness for mapped interactions are proportional to IntAct MI score, else they are colored in gray with constant thickness. Node rims are colored according to enriched processes; (C) Dot plot showing top 15 significantly enriched (FDR < 0.01) categories from the LRRK2-MLi-2 interactome. Values on the x axis display the ratio of genes of the term participating to the network over the total number of genes in that category. The ratio of the dot is proportional to the number of genes of that category in the network, while the node color is the darker the more significant is the category enrichment. Only categories with at least 5 genes have been displayed. Representative, enriched categories have been colored with the same color scheme as in panel C. (D) Scheme highlighting the function of centriolar satellites and the core protein PCM1 in ciliation (Hori & Toda, 2017, modified) (E) BioID analysis following PCM1 knockdown and MLi-2 treatment. Shown are eluates obtained after streptavidin-based enrichment of biotinylated proteins. (F) Violin plot reporting the Jaccard Score distribution in different data sets. P-values have been computed with a two-sided Mann-Whitney test with Bonferroni correction (* *P* < 0.05; P = 0.0499 for T-test BioID vs MLi-2 BioID; comparison between 188 BioID, 431 T-test BioID, 36 RAB29 BioID and 9 MLi-2 BioID independent samples; only P-values < 0.05 are shown). Violin plots show the median as the white central dot and first and third quartiles as bounds of the thicker black line (box); the whiskers extend to the last data point within 1.5 times the interquartile range (IQR) from the box’s boundaries; (G) Volcano-plot (+/-RAB29): mT-LRRK2 proximity proteome following RAB29 co-expression (moderated FDR-based T-test, P = 0.05; S0 = 0.1, N = 9). RAB29 redirects LRRK2 to lysosomal PPI networks; (H) Stringified network of the RAB29 over-expression interactome from (G) mapped to IntAct known interactions. Edges’ color and thickness for mapped interactions are proportional to IntAct MI score, else they are colored in gray with constant thickness. Node rims are colored according to enriched processes; (I) Dot plot showing top 15 significantly enriched (FDR < 0.01) categories from the LRRK2-RAB29 over-expression interactome. Values on the x axis display the ratio of genes of the term participating to the network over the total number of genes in that category. The ratio of the dot is proportional to the number of genes of that category in the network, while the node color is the darker the more significant is the category enrichment. Only categories with at least 5 genes have been displayed. Representative, enriched categories have been colored with the same color scheme as in panel H.

**Table 2:** Enriched proteins in the LRRK2 proximity proteome upon MLi-2 treatment (FDR-controlled T-test).

| Accession | Protein name | Gene name | P-value | Log2 T-test difference |
| --- | --- | --- | --- | --- |
| P81605 | Dermcidin | DCD | 2.91844 | 3.3 |
| Q9UPN4 | Centrosomal protein of 131 kDa | CEP131 | 4.15494 | 2.2 |
| Q49A88 | Coiled-coil domain-containing protein 14 | CCDC14 | 5.39373 | 2.1 |
| Q9Y2D8 | Afadin- and alpha-actinin-binding protein | SSX2IP | 2.98866 | 2.1 |
| O75665 | Oral-facial-digital syndrome 1 protein | OFD1 | 7.92350 | 2.0 |
| Q15154 | Pericentriolar material 1 protein | PCM1 | 6.62987 | 1.8 |
| Q9UHB6 | LIM domain and actin-binding protein 1 | LIMA1 | 2.96451 | 1.7 |
| Q8WXW3 | Progesterone-induced-blocking factor 1 | PIBF1 | 4.62801 | 1.5 |
| Q6WCQ1 | Myosin phosphatase Rho-interacting protein | MPRIP | 7.40967 | 1.5 |
| Q14596 | Next to BRCA1 gene 1 protein | NBR1 | 3.34347 | 1.3 |
| Q5TB80 | Centrosomal protein of 162 kDa | CEP162 | 3.46643 | 1.3 |
| Q8WWI1 | LIM domain only protein 7 | LMO7 | 5.84664 | 1.3 |

To functionally validate the interaction between LRRK2 and centriolar satellites, we first confirmed that LRRK2 associates with key components of these membrane-less organelles, including the central scaffold protein PCM1 and the centrosome maturation factor SSX2IP (Barenz *et al*, 2013). Both proteins were enriched in LRRK2 proximity proteomes following biotin treatment, with their interaction further enhanced by the LRRK2 inhibitor MLi-2. To further substantiate LRRK2’s association with centriolar satellites, we employed RNA interference to deplete PCM1, a strategy known to disrupt satellite integrity (Dammermann & Merdes, 2002). As expected, PCM1 depletion led to its absence from the LRRK2 proximity proteome and also abolished the association between LRRK2 and SSX2IP. Remarkably, SSX2IP was entirely absent from LRRK2 proteomes regardless of MLi-2 treatment, indicating that LRRK2’s interaction with SSX2IP is dependent on intact centriolar satellites. Finally, we asked whether the MLi-2–induced dephosphorylation of serine 935—located near the LRRK2 hinge helix—is affected by satellite disruption. However, PCM1 knock-down did not influence serine 935 dephosphorylation, suggesting that this post-translational modification occurs independently of centriolar satellite integrity (Figure 6D, Appendix Figure S6C).

### RAB29 co-expression directs LRRK2 to well established lysosomal protein networks

It has been previously shown that LRRK2 interacts with the RAB29/RAB32/RAB38 family of small G-proteins and subsequently locates to post-Golgi endomembranes (Beilina *et al*, 2014; McGrath *et al*, 2021; Vides *et al*, 2022; Waschbusch *et al*, 2014). Furthermore, RAB29 not only recruits LRRK2 to lysosomal membranes but also induces its tetramerization (Zhu *et al*, 2023). For this reason, we performed the BioID experiment following RAB29 co-expression to determine PPI networks associated with the activated oligomeric state of LRRK2. In addition to mT-LRRK2, either RAB29 or an empty vector were co-expressed. The resulting differential proximity proteome showed differentially enriched proteins only for the condition where RAB29 was co-expressed, demonstrating that a small portion of LRRK2 was recruited by RAB29 with lesser impact on the total proximity proteome (Figure 6G). While we did not observe any significant difference in terms of co-evolution of the RAB29 differential proteome with respect to the BioID one (Figure 6F), we found significant enrichment in proteins involved in vesicular transport, including the post-Golgi/ lysosomal compartment as revealed by an LFQ analysis combined with an FDR-controlled T-test (Figure 6H, I). In particular, known LRRK2 effectors, namely RAB8A (Steger *et al*, 2017) and SPAG9 (Boecker & Holzbaur, 2021; Bonet-Ponce *et al*, 2020; Kluss *et al*, 2022a) were among the enriched proteins (Figure 6G, H). Also, VAMP7, which is part of a SNARE complex mediating autophagosome-lysosome fusion (Jian *et al*, 2024), was among the significantly enriched proteins (Figure 6G, H). VAMP7-LRRK2 physical interaction has recently been described (Filippini *et al*, 2023). The complete list of significantly enriched proteins is shown in Table 3. Additional information is provided in Dataset EV1.

**Table 3:** Enriched proteins in the RAB29-dependent LRRK2 proximity proteome (FDR-controlled T-test).

| Accession | Protein name | Gene name | P-value<br>(-log10) | Log2 T-test<br>difference |
| --- | --- | --- | --- | --- |
| O14966 | Ras-related protein Rab-7L1 | RAB29 | 9,72545 | 10,5 |
| Q8IZA0 | Dyslexia-associated protein KIAA0319-like protein | KIAA0319L | 5,00209 | 4,9 |
| Q6WKZ4 | Rab11 family-interacting protein 1 | RAB11FIP1 | 4,56815 | 4,6 |
| O15126 | Secretory carrier-associated membrane protein 1 | SCAMP1 | 4,09385 | 3,5 |
| Q9UIQ6 | Leucyl-cystinyl aminopeptidase;Leucyl-cystinyl<br>aminopeptidase, pregnancy serum form | LNPEP | 4,33552 | 3,4 |
| Q9H2J7 | Sodium-dependent neutral amino acid transporter<br>B(0)AT2 | SLC6A15 | 5,06972 | 2,9 |
| Q96QD8 | Sodium-coupled neutral amino acid transporter 2 | SLC38A2 | 2,89271 | 2,9 |
| Q13586 | Stromal interaction molecule 1 | STIM1 | 3,12115 | 2,8 |
| P98194 | Calcium-transporting ATPase type 2C member 1 | ATP2C1 | 5,00347 | 2,6 |
| Q9H2H9 | Sodium-coupled neutral amino acid transporter 1 | SLC38A1 | 2,37723 | 2,6 |
| O75976 | Carboxypeptidase D | CPD | 3,20952 | 2,6 |
| Q14126 | Desmoglein-2 | DSG2 | 4,31421 | 2,5 |
| P51809 | Vesicle-associated membrane protein 7 | VAMP7 | 4,02209 | 2,5 |
| Q8NDI1 | EH domain-binding protein 1 | EHBP1 | 2,37346 | 2,5 |
| Q8TBA6 | Golgin subfamily A member 5 | GOLGA5 | 3,0259 | 2,4 |
| Q9BXF6 | Rab11 family-interacting protein 5 | RAB11FIP5 | 2,9101 | 2,4 |
| Q7L804 | Rab11 family-interacting protein 2 | RAB11FIP2 | 3,33154 | 2,2 |
| Q6IAA8 | Ragulator complex protein LAMTOR1 | LAMTOR1 | 3,23604 | 2,1 |
| P61006 | Ras-related protein Rab-8A | RAB8A | 3,56486 | 2,1 |
| P11717 | Cation-independent mannose-6-phosphate<br>receptor | IGF2R | 5,71543 | 2,0 |
| Q13190 | Syntaxin-5 | STX5 | 3,88884 | 2,0 |
| Q15836;P63027 | Vesicle-associated membrane protein 3;Vesicle-associated membrane protein 2 | VAMP3;VAMP2 | 3,67019 | 2,0 |
| Q92692 | Nectin-2 | PVRL2 | 3,06334 | 2,0 |
| Q6NXT4 | Zinc transporter 6 | SLC30A6 | 3,47208 | 1,8 |
| Q96A65 | Exocyst complex component 4 | EXOC4 | 2,51947 | 1,7 |
| Q14141 | Septin-6 | Sept6 | 3,87782 | 1,7 |
| Q04656 | Copper-transporting ATPase 1 | ATP7A | 2,77415 | 1,7 |
| Q7Z2W4 | Zinc finger CCCH-type antiviral protein 1 | ZC3HAV1 | 2,62567 | 1,6 |
| P78310 | Coxsackievirus and adenovirus receptor | CXADR | 2,87956 | 1,4 |
| P08195 | 4F2 cell-surface antigen heavy chain | SLC3A2 | 2,43497 | 1,4 |
| P02786 | Transferrin receptor protein 1;Transferrin<br>receptor protein 1, serum form | TFRC | 3,21618 | 1,3 |
| Q07960 | Rho GTPase-activating protein 1 | ARHGAP1 | 2,65929 | 1,3 |
| O60271 | C-Jun-amino-terminal kinase-interacting protein 4 | SPAG9 | 3,80826 | 1,3 |
| P42167 | Lamina-associated polypeptide 2, isoforms<br>beta/gamma;Thymopoietin;Thymopentin | TMPO | 3,54449 | 1,2 |
| Q9BSJ8 | Extended synaptotagmin-1 | ESYT1 | 3,59064 | 1,0 |
| Q14739 | Lamin-B receptor | LBR | 3,12029 | 0,8 |

### MLi-2 and RAB29 differently modulate the shape of the LRRK2 interactome

We constructed 3D models of binary complexes between LRRK2 and its partners from the MLi-2–and RAB29-dependent interactomes, and examined both the complex interfaces and the conformational states of LRRK2 (Vides *et al*, 2022).

Interface-based clustering suggests that MLi-2–dependent interactors engage with LRRK2 through specific domain regions, particularly the ones involving terminal domains (Figure EV1 A), while they are predicted to engage little with the catalytic domains, particularly the kinase, which is consistent with the MLi-2 inhibitor function. In the RAB29-dependent LRRK2 interactome, we see instead a more variable pattern of engaged interfaces. In particular, predictions for this interactome revealed a cluster of proteins interacting with LRRK2 through the ROC-COR-KIN domains, including the RAB8A substrate (Figure EV1B), suggesting that in the context of RAB29 overexpression several partners might bind in a substrate-like manner. Notably, this interaction modality is absent in the predicted binary complexes of the MLi-2–bound interactome, confirming that the inhibitor prevents LRRK2 from binding substrate-like partners. Moreover, we further validated the capability of the distogram-based interface-fingerprint to discriminate functionally relevant LRRK2 interactors. We predicted 3D complexes between LRRK2 and known interacting Rabs and found that Rab substrates clustered separately from other interacting Rabs with distinct functions, such as RAB29 and RAB32 (Figure EV1B).

From a conformational perspective, we pooled the RAB29- and MLi-2–dependent interactomes and clustered LRRK2 structures based on RMSD, which likewise revealed distinct locked and unlocked clusters (Figure 7A). Interestingly, Rab8a acting downstream of LRRK2 and Rab29 acting upstream of LRRK2 engage with LRRK2 in two different conformations: While Rab8a engages with the unlocked state (Figure 7B), structural modelling suggests that Rab29 engages with the locked state (Figure 7C). Interactors from the RAB29-dependent interactome are predicted to bind LRRK2 in both conformations, with a particular enrichment in the unlocked state. Conversely, MLi-2–dependent interactors were significantly enriched in the locked LRRK2 conformation (Fisher’s exact test, P = 0.023), further suggesting that the inhibitor-bound state favors engagement with partners that, unlike substrates, do not require interaction with the kinase domain (Figure 7A). Also for these interactors, the locked conformation has a significantly lower RMSD (P = 3.85E-6; Figure 7D) and a tendency towards higher co-evolution (Figure 7E). Notably, enrichment analysis of interaction partners grouped by conformational cluster revealed even stronger cluster-specific associations, with only a single process shared between partners in the locked and unlocked conformations (Figure 7F, G; Dataset EV2E).

**Figure 7:**
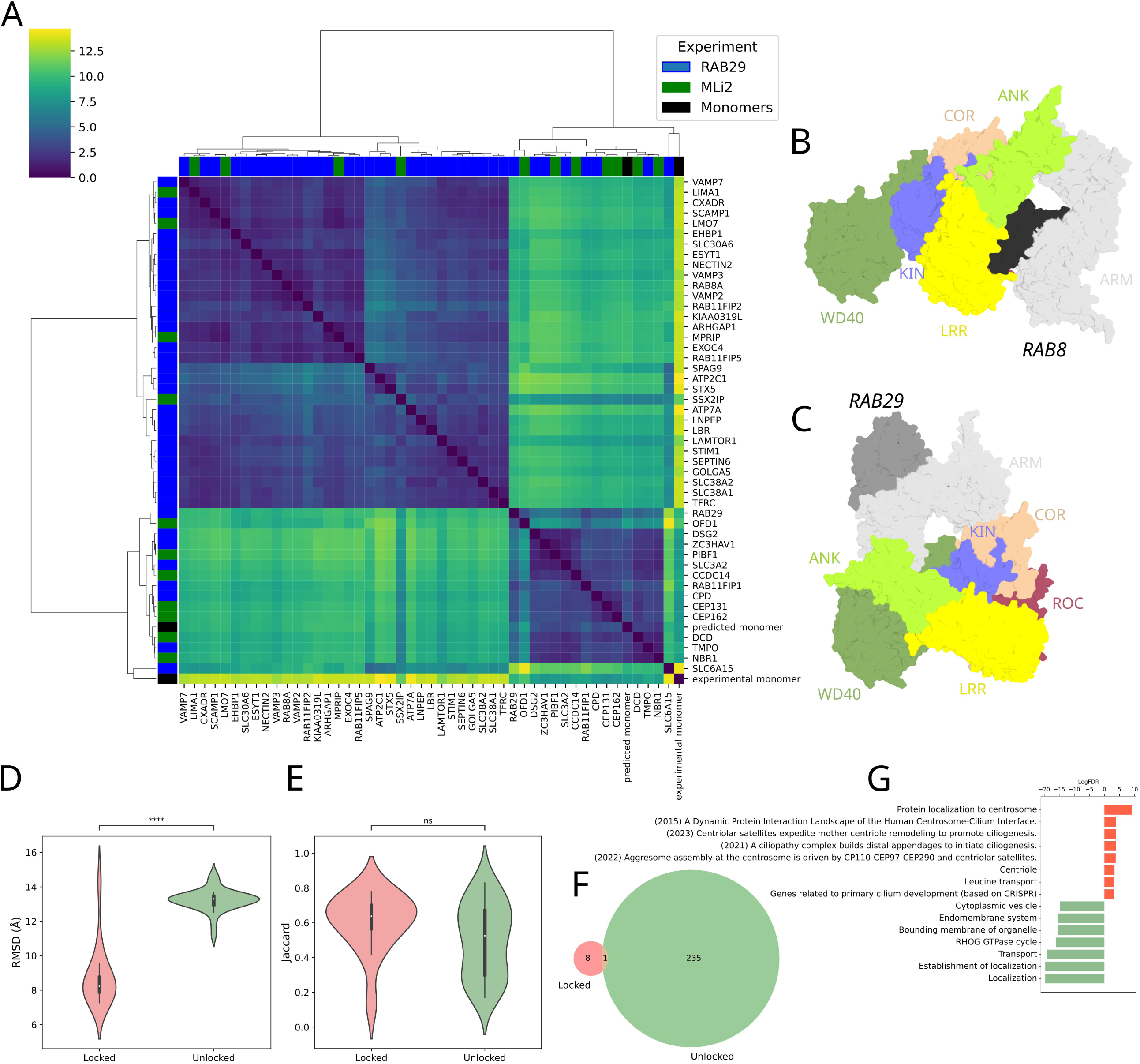
AF-structural modeling of MLi-2 and Rab29 interactomes. (A) Hierarchical clustering of pairwise RMSD of LRRK2 chain in the different complexes; B) LRRK2-RAB8A predicted complex; (C) LRRK2-RAB29 predicted complex; (D) violin plot of the distribution of RMSD calculated with respect to the reference LRRK2 experimental structure (PDB 7LHW) for the locked and unlocked clusters. Color scheme as in 4C (P = 3.8E-6; comparison between 14 locked and 32 unlocked independent samples); (E) distribution of the Jaccard score for the locked and unlocked clusters (P = 0.14; comparison between 14 locked and 32 unlocked independent samples). Color scheme as in 4C. P-values in violin plots in (D) and (E) have been computed with a two-sided Mann-Whitney test (* *P* < 0.05, ** *P* < 0.01, *** *P* < 0.001, **** *P* < 0.0001). Violin plots show the median as the white central dot and first and third quartiles as bounds of the thicker black line (box); the whiskers extend to the last data point within 1.5 times the interquartile range (IQR) from the box’s boundaries; (F) Venn diagram of the enriched terms in the locked and unlocked clusters. Color scheme as in 4C; (G) Top exclusively enriched (FDR < 0.01) terms in the locked and unlocked clusters. Color scheme as in 4C.

## Discussion

The systematic analysis of the interactomes of disease-associated proteins is a powerful tool to dissect molecular networks and to identify pathways relevant for the molecular pathophysiology, for example demonstrated by the comprehensive interactomic network underlying the assembly and homeostasis of primary cilia (Boldt *et al*, 2016). Given its complex domain structure and tight regulation, LRRK2 is most likely representing a scaffold protein orchestrating distinct signaling complexes, which makes the analysis of its PPI networks an attractive approach to dissect context-specific signaling pathways (Gloeckner & Porras, 2020). While a wealth of data is available for affinity-based identification of protein-protein interactions, which are biased toward stable complexes, there is an unmet need for the dissection of less stable and transient PPIs. The rapid development of biotin-based proximity labelling technologies has made it possible to explore previously inaccessible regions of the interactome (Qin *et al*, 2021). Consequently, APEX and BioID have emerged as widely used techniques for dissecting protein-protein interaction (PPI) networks (Li *et al*, 2017). For LRRK2, the first APEX2 dataset only recently became available (Bonet-Ponce *et al*, 2020). In the present study, we systematically investigated the proximity interactome of LRRK2 using three different BioID tags—BioID1, BioID2 and miniTurbo. The resulting data showed the expected overlap with previously reported interactors, while also uncovering numerous novel candidates. However, several anticipated interactors, including Rab proteins, were not detected in the initial BioID experiments. Nevertheless, Rab8a was identified together with Rab29 upon co-expression of Rab29, which may reflect conformational constraints that reduce accessibility for BioID-mediated biotinylation. Consistent with these observations, a recent comparison of BioID and co-immunoprecipitation in the context of the nuclear pore complex demonstrated that the two approaches provide functionally relevant yet complementary interaction profiles (Moreira *et al*, 2023).

To identify direct LRRK2 interactors with high confidence from the proximity proteomes, we applied a newly developed bioinformatic pipeline relying on co-evolution, co-expression and protein domain architecture to predict interaction type (preprint: Miglionico *et al*, 2024). This pipeline allowed the identification of a group of proteins within the proximity proteome showing a high degree of co-evolution with LRRK2 which is highly enriched in proteins associated with the centrosome. This is particularly interesting as pathogenic LRRK2 has previously been shown to lead to a centrosome cohesion defect as well as to a shortening of primary cilia (Dhekne *et al*, 2018; Khan *et al*, 2024; Lara Ordonez *et al*, 2022). Furthermore, defects in primary cilia emerge as a new pathophysiological scheme connecting different genes involved in mendelian forms of Parkinson’s disease (Mohd Rafiq *et al*, 2024; Schmidt *et al*, 2022). Strikingly, within the cluster of proteins showing the highest co-evolution with LRRK2 (Cluster 2), the top-ranked interactor was the deubiquitinase CYLD, a key effector in primary cilia biology. Preliminary data further indicate that CYLD co-expression increases the levels of the pathogenic LRRK2 Y1699C variant, which exhibits lower steady-state protein levels than wild-type LRRK2. In fact, this DUB has been shown to control apical docking of basal bodies in ciliated epithelial cells (Eguether *et al*, 2014). It also regulates centrosomal satellites proteostasis by counteracting the E3 ubiquitin ligase MIB1 (Douanne *et al*, 2019). CYLD has also been shown to deubiquitinate Cep70 and to inactivate HDAC6 thereby mediating ciliogenesis (Yang *et al*, 2014). Interestingly, a missense mutation has recently been shown to be a causative gene for FTD-ALS (Dobson-Stone *et al*, 2020). Most importantly, as the deubiquitinase CYLD is part of the PINK1/Parkin pathway to regulate cellular levels of PARIS (ZNF746), it is a critical regulator of mitochondrial biogenesis thereby acting as a negative regulator of dopamine neuron survival in Parkinson’s disease. In fact, CYLD inhibition has been shown to ameliorate mitochondrial pathologies in cellular PD models which makes CYLD a promising target for therapeutic intervention for PD and other neurodegenerative diseases (Pirooznia *et al*, 2022). The E3 ligase Parkin preserves mitochondrial homeostasis via the PARIS/PGC-1a axis (Stevens *et al*, 2015; Zheng *et al*, 2017). Noteworthy, the PD risk variant LRRK2 G2385R has recently been shown to alter mitochondrial biogenesis via the PGC-1a pathway, leading to reduced PGC-1a levels similarly to a Parkin knock-out in DA neurons (Kumar *et al*, 2020; Xue *et al*, 2023). Parkin-deficient DA neurons show increased PARIS and decreased PGC-1a levels (Kumar *et al*, 2020). In conclusion, this indicates that LRRK2/CYLD signaling, at one hand, and Parkin signaling at the other, might converge on PARIS.

The second most co-evolved interactor found in Cluster 2 was the cilia-associated protein CEP85, which plays a role in centrosome disjunction (Chen *et al*, 2019). This is of particular interest as pathogenic LRRK2 variants lead to a centrosomal cohesion phenotype in patient-derived cells (Lara Ordonez *et al*, 2019). Notably, the cohesion phenotype is also seen in EBV-transformed lymphoblastoid cell lines from iPD patients which can be reverted by LRRK2 kinase inhibition (Naaldijk *et al*, 2024). Importantly, Cluster 2 also contains the autophagy cargo protein p62/sequestosome-1. This protein has previously been identified as LRRK2 interactor and substrate (Kalogeropulou *et al*, 2018; Park *et al*, 2016). SQSTM1 is also linked to PD pathophysiology via the Pink1/Parkin pathway (Shin *et al*, 2020). In addition, Cluster 2 points towards other pathways which have been previously linked to PD pathology. In particular, SNAP47 is part of the STX17-SNAP47-VAMP7/8 pathway, which has been shown to be involved in selective autophagy – a process not only essential for the homeostasis of primary cilia but also for mitophagy (Jian *et al*, 2024). Phospho-ubiquitin chains generated by PINK1/Parkin activity can recruit autophagy adaptors including the ubiquitin adaptor SQSTM1/p62 linking the PINK1/Parkin-dependent pathway to mitophagy (Geisler *et al*, 2010). All three proteins participate in the recruitment of the autophagy machinery to aged or dysfunctional mitochondria, which are eventually enclosed inside an autophagosome and subjected to the lysosome for degradation (Wilhelm *et al*, 2022).

Interestingly, also the kinesin motor protein and PX-domain containing protein KIF16B was among the top hits which is involved in vesicle trafficking and receptor recycling. It binds to PdIns(3)P by its unique PX domain and promotes the circulating cargo movement from early endosomes to the plasma membrane (Li *et al*, 2020). Interestingly, it has been found to be associated with lipid droplets (Bersuker *et al*, 2018). In turn, increased RAB8a phosphorylation by pathogenic LRRK2 variants, has been shown to promote lipid storage (Yu *et al*, 2018), indicating that the functional interaction of LRRK2 and KIF16B might contribute to alterations in the neuronal lipid homeostasis associated with PD pathology (Hallett *et al*, 2019).

Furthermore, conformational changes induced by MLi-2 binding—leading to rapid and nearly complete dephosphorylation of LRRK2 within a disordered domain downstream of the hinge helix—position LRRK2 in proximity to a molecular network centered around PCM1, OFD1, and CEP131, strongly supporting a link between LRRK2 and centriolar satellites. Strikingly, the proteins interacting with LRRK2 in the presence of MLi-2 displayed the highest degree of co-evolution, suggesting that the connection between LRRK2 and centriolar satellites is highly conserved. The conformational changes induced by binding of type I inhibitors might result in a phase transition of LRRK2 into centriolar satellites preventing 14-3-3 binding and thereby shifting the equilibrium towards the dephosphorylated protein. A similar mechanism has been shown for CEP131, which is extracted from centriolar satellites by phosphorylation and subsequent binding of 14-3-3 (Tollenaere *et al*, 2015). Centriolar satellites not only play an essential role in the biogenesis of primary cilia (Hall *et al*, 2023) but also play a role in the degradation of proteins by driving the assembly of aggresomes (Prosser *et al*, 2022). In fact, also LRRK2 has been shown to interfere with aggresome formation preceding autophagic clearance (Bang *et al*, 2016). Nevertheless, one caveat might be seen in the fact that LRRK2 inhibition with type I inhibitors induces its destabilization and degradation (Lobbestael *et al*, 2016).

Structural prediction of LRRK2 binary complexes with AF revealed distinct interfaces within each determined interactome. While several interactors preferentially engage with the catalytic core of the protein, including the ROC and KIN domains, others are more likely to interact on terminal domains (e.g. ARM or WD40). We predicted the interaction on the kinase domain for several interactors of the BioID interactome as well as the one obtained in the presence of RAB29 over-expression, including the known substrate RAB8A. The engagement in a substrate-like fashion with the catalytic core entails the opening of the LRRK2 structure, with the detachment of the LRR and rearrangement of the entire N-term domains. We previously showed that this conformation, that we similarly predicted for the LRRK2:RAB10 complex, is compatible with distance restraints obtained by mass spectrometric mapping of chemical crosslinking sites via XL-MS (Guaitoli *et al*, 2023). Moreover, when superimposed onto the active-like conformation of the most recent LRRK2 tetrameric structure, these predicted complexes fit well within the oligomer, without steric clashes between the modeled LRRK2 chain and the rest of the tetramer, further supporting their structural plausibility (Guaitoli et al, 2023). We speculate that partners engaging in this manner may represent putative substrates or function as regulators of kinase domain activity. Strikingly, such interaction interfaces were rarely observed for interactors enriched in the presence of the MLi-2 inhibitor, which preferentially bind the locked conformation of LRRK2.

In conclusion, combining proximity proteomics with co-evolutionary and structural information helped identify distinct molecular networks linking LRRK2 to centrosomal biology and primary cilia. The data may foster future functional analysis to further dissect these pathways to get a detailed picture of the molecular pathophysiology underlying the cilia phenotypes caused by genetic variants in different genes underlying familial forms of Parkinson’s disease.

## Methods

### Cloning and generation of the stable miniTurbo-LRRK2 cell line

For comparability, all BioID tags used in this study were generated using an identical cloning strategy, incorporating an N-terminal HA tag for detection and a 3×(GGGS) linker between the tag and the protein of interest, inserted C-terminally via Gateway cloning. The constructs have been ordered as synthetic genes (Eurofins) and subcloned via *Eco*RI and *Xho*I into pcDNA3.0 (Invitrogen) allowing the expression of fusion proteins in mammalian cells under the strong viral CMV promoter. Gateway Destination clones were generated by cloning a Gateway cassette (Invitrogen) into an *Eco*RV site at the 3’ end and in frame of the different BioID-tags [BioID (Roux *et al*, 2012; referred as BioID1), BioID2 (Kim *et al*, 2016), miniTurbo (Branon *et al*, 2018)]. For better and comparable detection of the fusion proteins, all tags contained an N-terminal HA-tag. The LRRK2 expression clones were subsequently generated by LR cloning (Invitrogen) based on ENTRY clones described earlier (Gloeckner *et al*, 2010). The CDS of human of RAB29 was ordered as synthetic gene strand (Eurofins) with flanking attB sites to allow its direct cloning into pDONR201 by BP cloning and subsequently cloned into an in-house (N)FLAG-HA Gateway destination clone by the LR reaction. The HEK293 cell line (ACC-305) stably expressing miniTurbo-LRRK2 was generated as previously described (Gloeckner *et al*, 2007). Briefly, single colonies have been selected and tested after transient transfection and selection via G418 (Sigma). The Flag-HA-CYLD vector was kindly provided by Wade Harper (plasmid #22544, Addgene) (Sowa *et al*, 2009), the pCMV poly HA-Ubiquitin construct was kindly provided by D. Bohmann (Treier *et al*, 1994) and the pRK5-HA-Ubiquitin-K63 vector was kindly provided by T. Dawson (plasmid #17606, Addgene) (Lim *et al*, 2005).

### Cell culture and proximity labelling

HEK293T (ACC-635) or HEK293 stably expressing miniTurbo-LRRK2 were grown in 14cm dishes in full Dulbecco’s modified medium (DMEM) supplemented with penicillin streptomycin and if not mentioned otherwise with 10% FBS at 37°C in 5% CO2. The cells were grown for 8 h to 1 day reaching a density of 50-80% until transfection with a self-made PEI (Polyethylenimine) reagent (Guaitoli *et al*, 2016). For each condition, 4–6 plates were used. If not stated otherwise, for biotinylation, cells have been incubated with 50 µM biotin (Sigma) in serum-free medium for up to 16 hours. In case of the mini turbo tag, cells have been incubated with 50 µM biotin for 2 h after o/n starvation. Starvation was used to synchronize the cells and, in light of the physiological functions of LRRK2, to enrich for post-mitotic states in which its centrosomal and ciliary roles are more prominent. For the determination of the LRRK2-inhibitor dependent proximity proteomes, a HEK293 line stably expressing mT-LRRK2 was either incubated with 1 µM MLi-2, 1 µM GZD-824 or DMSO (vehicle) 1h prior and during 2 h biotin incubation. For the analysis of RAB29-dependent proximity proteomes, mT-LRRK2 was either co-expressed with HA-RAB29 or an empty pcDNA3.0 vector. See also the point-to-point protocol deposited on Zenodo (DOI: 10.5281/zenodo.17045153).

### Affinity purification of biotinylated proteins

For cell lysis, cell culture dishes were transferred to ice and culture medium was removed. The cells were rinsed two times with 5ml ice-cold PBS. The cells were covered with 1ml lysis buffer [1% Nonidet P-40 Substitute (Merck), protease inhibitor cocktail (Roche), PhosSTOP (Roche) in TBS (30 mM Tris-HCl, pH7.4 and 150 mM NaCl)] per 14 cm dish and detached with a cell scraper. The cell suspension was collected and incubated for 20-40 min on an end-over-end shaker at 4°C. Lysates were cleared by centrifugation (15,000 x g for 10 min at 4°C). The protein concentration of cleared lysates was determined photometrically using a nanodrop (Thermo-Fisher Scientific) or with the BCA assay (Thermo-Fisher Scientific) following the manufacturer’s instructions. Biotinylated proteins were purified by affinity enrichment using either Strep-Tactin Superflow resin (IBA) or MagStrep Strep-Tactin beads (IBA) in case of small-scale enrichments. To this end, the combined lysates for each condition were incubated with an adjusted resin volume of 25 µl packed beads per 14cm cell culture dish. The resin was equilibrated by washing 3-4x with 500 µl lysis buffer. The lysates were incubated with the resin overnight at 4°C under constant agitation. After incubation, the resin was sedimented by centrifugation at 1,000 x g for 3 min at 4°C. The supernatant was removed, leaving approximately 300 µl to resuspend the resin. The resin suspension was transferred to microSpin columns (Cytiva) and washed three times with 500 µl lysis buffer, followed by two washes with 500 µl 1x TBS (centrifugation for 5 s at 100 x g). Biotinylated proteins were eluted by incubation with 300-400 µl desthiobiotin elution buffer (2.5M, IBA). The desthiobiotin eluates were precipitated by chloroform/ methanol and directly subjected to tryptic proteolysis following standard protocols (Gloeckner *et al*, 2009) and stored at −80°C. The BioID protocol was further optimized during the course of the study. A detailed step-by-step protocol has been deposited on Zenodo (DOI: 10.5281/zenodo.17045153). This protocol includes an additional SDS extraction step after cell lysis and removal of nuclei and was used in particular for the inhibitor study.

### CYLD co-expression experiment

For the analysis of CYLD co-expression, cells were co-transfected with either Strep/FLAG (SF-TAP)-tagged LRRK2 WT or LRRK2 Y1699C together with CYLD. As negative control, CYLD plasmid was replaced by pCDNA3 empty vector, to maintain the plasmid concentration constant among the different conditions. To test the influence of defined ubiquitin linkage types, ubiquitin—either in WT form (pCMV poly HA-Ubiquitin) or K63 (pRK5-HA-Ubiquitin-K63)—was co-expressed. For experiments relying on endogenous ubiquitin, the ubiquitin expression plasmids were replaced by the pcDNA3 vector. The experiment was performed in 10 cm culture dishes. Cells were seeded at a density of 3 to 4 million cells per dish and transfected the next day using home-made PEI transfection reagent (cell confluency 30-40%). A total of 3.2 µg of plasmid DNA was used for each plate. The plasmids ratios were LRRK2:CYLD:Ubiquitin 1:3:1. Cells were lysed 48 h post-transfection (cell confluency approx. 80%) in 500 µl lysis buffer (see above) and incubated for 45 min at 4°C under agitation. Cell debris was removed by centrifugation at 1,000 × g for 10 min at 4°C. To extract membrane-bound prenylated proteins (e.g., Rab proteins), pre-cleared lysates were supplemented with SDS (55 µl of a 10% stock solution) to a final concentration of 1% and incubated for 15 min at 4°C under agitation. Lysates were subsequently clarified by centrifugation at 20,000 × g for 10 min at room temperature. Protein concentration was determined by BCA assay (Pierce), and LRRK2 levels were assessed by capillary Western blotting (see below).

### siRNA knockdown

24h after splitting, cells were transfected with RNAiMAX (Life Technologies) following manufacturer’s instructions. The following siRNAs were used (Silencer Select®, Life Technologies): PCM1 (#s10128, CAAAGACUCCACAUACGUUtt), Negative Control No. 2 (#4390846). 360 pmol of siRNA were used for each technical replicate. Cells were kept in culture for another 48h before proceeding with treatment with 10µM MLi-2 or DMSO. Cells were lysed as described above. Three technical replicates for each condition (PCM1 or scrambled, MLi-2 or DMSO) were used. The effects of PCM1 knockdown were evaluated via SDS-PAGE and Western Blot.

### SDS PAGE and Western Blots

For functional evaluation of the bait proteins and to determine the biotinylation efficiency, samples were subjected to SDS PAGE and analyzed by Western-blotting on PVDF membranes. To assess blotting efficiency and equal loading, the membranes were stained with Ponceau S. Prior to blocking of the membranes in 5% nonfat dry milk powder (BioRad) or 1-5% BSA (Carl Roth) in TBS with 0.1% Tween 20 (TBST) for 1-3h at RT, membranes were de-stained in TBST. Membranes were afterwards incubated with primary antibody solutions overnight at 4°C under mild agitation. The following primary antibodies were used: rat α-LRRK2 mAb (24D8 clone, hybridoma supernatant, generated in-house (Carrion *et al*, 2017)) 1:1000 in 5% milk in TBST), rabbit α-pS935-LRRK2 mAb (Abcam, ab133450; 1:100,000 in 5% BSA in TBST), rabbit α-RAB10 mAb (Abcam, ab181367; 1:5,000 in 5% milk in TBST), rabbit α-pT73-RAB10 mAb (Abcam, ab230261; 1:1,000 in 5% BSA in TBST), rat α-HA-tag (Roche, 3F10 clone, Rat 1gG1; 1:1,000 in 5% milk in TBST), mouse α-FLAG-tag (Sigma, F1804-200UG, 1:2,000 in 5% milk in TBST), rabbit α-PCM1 (Proteintech, 19856-1-AP, 1:24,000 in 5% milk in TBST), mouse α-CYLD (Proteintech, 66858-1-Ig, 1:1,500 in 5% milk in TBST), rabbit α-SSX2IP (Thermo Fisher Scientific, PA5-31495, 1:1,000 in 5% milk in TBST). Membranes were washed 5x 5min in TBST prior to incubation with the HRP-conjugated secondary antibodies α-rat (Jackson Immuno-Research, 112-036-062; 1:7,500), α-rabbit (Jackson Immuno-Research, 111-036-045; 1:7,500), and α-mouse (Jackson Immuno-Research, 115-036-062; 1:7,500), respectively. For the detection of biotinylated proteins, membranes were incubated with Pierce™ High Sensitivity Streptavidin-HRP (Thermo Fisher Scientific, 1:20,000 in 1% BSA in TBST) overnight at 4°C and washed the next day 5x 5min in TBST. Blots were developed with Pierce ECL Plus Western Blotting Substrate following the manufacturer’s protocol (Thermo Fisher Scientific) and exposed to Amersham Hyperfilm™ ECL (Cytiva). For quantification of LRRK2 in presence of CYLD, the JESS capillary Western system (Biotechne) was used. Separation was performed on a 25-capillary array (12-230kDa Fluorescence Separation; SM-FL004-1) LRRK2 was detected by 24D8; 1:25/ rabbit anti-rat DyLight-633; 1:50 (Biotechne NBP1-76225) in milk diluent. b-actin was detected by anti-b-actin; 1:500 (R&D Systems MAB8929)/ anti-mouse-NIR (Biotechne 043-821); 1:20; in milk diluent. Detection for both proteins was done in the NIR channel. Data analysis was performed using vendor software (Compass for SW, v7.1.0) and custom Python scripts; graphical representations were generated with the Python library Seaborn. Unpaired two-tailed t-tests were used to compare LRRK2 levels normalized by b-actin.

### Mass spectrometry

In solution proteolysis was performed as previously described (Gloeckner *et al*, 2009). Briefly, chloroform/ methanol precipitates were dissolved in 50 mM ammonium bicarbonate pH8.0. Cysteines were reduced and alkylated by treatment with 100 mM DTT and 300mM Iodoacetamide, respectively following the incubation with trypsin (Sigma-Aldrich) overnight at 37°C. Samples were desalted by micropipette C-18 solid-phase extraction tips (StageTip, Thermo Fisher Scientific) prior to LC-MSMS analysis. Peptides were subsequently analyzed by LC-MSMS using a nanoflow HPLC system (Ultimate 3000 RSLC; Thermo-Fisher Scientific) coupled to an Orbitrap Q-Exactive plus (Thermo Fisher Scientific) tandem mass spectrometer. Peptides were separated by reversed C-18 chromatography and 120-minute gradients. MS1 spectra were acquired in the Orbitrap at 70K resolution. After selection of the 10 most intense precursor ions from the MS1 scans for HCD fragmentation (Top 10 method), MS2 spectra were acquired at 17.5K. RAW MS files were analyzed with Maxquant (version 2.0.3.1), enabling label-free-quantification (LFQ with no normalization) and “match between runs” (Cox *et al*, 2014). For the embedded Andromeda search engine, trypsin has been selected as enzyme and carbamylation was set as fixed modification N-terminal acetylation was set as variable modification. The search was performed against the human subset of the Uniprot/Swissprot database (release 2021_04, 20375 entries). The MaxQuant output was further processed by Perseus and SaintExpress.

### Perseus post-processing

For post-analysis, the protein groups were further processed by Perseus (v. 1.6.7). LFQ intensities of the individual were assigned to the conditions and potential contaminants, reverse hits as well as IDs only identified by site were filtered out. For further analysis, only IDs with a maximum number of missing values of n-1 in at least one condition were considered. Remaining missing values were replaced by group-wise imputation based on a normal distribution (log2 values) around the detection limit (width=0.3; down-shift=1.8). Intensities of the individual experiments were normalized by their medians and significantly enriched proteins (condition vs control) were determined by a moderated permutation-based T-test (FDR = 0.05, S0 = 0.1) as implemented in Perseus (Two-sample test).

### SaintExpress post-processing

We followed a previously described protocol (Liu *et al*, 2020; see also http://proteomics.fi) to select the most confident interactors using the statistical framework of SAINTExpress (Teo *et al*, 2014), as well as a reference knowledgebase for contaminants (i.e. CRAPome (Mellacheruvu *et al*, 2013)), to filter out most likely false negative interactors. In particular, we generated the “Bait”, “Prey” and “Interaction” files to be subjected to the analysis pipeline from the MaxQuant “*proteinGroups.txt*” file by employing the following columns: “Bait” file (IP name = EV_* or LRRK2_*, bait name = LRRK2, indicator for test and negative control= C = EV_*, T = LRRK2_*); “Interaction” file (IP name = EV_*/LRRK_*, bait name = LRRK2, prey name = interaction partner gene symbol, spectral count or intensity values = LFQ intensity); “Prey” file (prey (protein)name = interaction partner Uniprot accession, prey protein length=Sequence length, prey gene name = gene symbol). The proteins that were not classified through the CRAPome filter, were retained together with those passing the filter.

We unified the interactor list from the three BioID experiments. Finally, we retained a list of 208 interactors which passed the two filters. Only 45 proteins in the filtered interactor list are already present in IntAct.

### Graphical representation and network annotation

PPI networks were analyzed by Cytoscape version 3.8.2 (Shannon *et al*, 2003) and were mapped to available interaction from IntAct using the Cytoscape IntAct app (Ragueneau *et al*, 2021). Networks were annotated with STRINGDB terms via the ‘Stringify’ function of the StringApp (Doncheva *et al*, 2019) and functional enrichment retrieved via the same app. We annotated the networks with the AutoAnnotate plug-in (Kucera *et al*, 2016), using the GLay clustering method, and generated the cluster labels with the Adjacent Words option of the WordCloud algorithm, based on significantly enriched GO Biological Process and Cellular Component terms associated with the genes in each network. The Venn diagram shown in Figure 1A has been generated with JVenn provided by the ProteoRE server (Galaxy Version 2021.05.12) (Galaxy, 2024).

### Co-evolution based PPI stratification

Co-evolution of two proteins A and B was measured as the Jaccard similarity coefficient of the set of genomes containing an orthologous sequence of A and the set of genomes containing an orthologous sequence of B. Orthologous sequences were collected from the Orthologous Matrix (OMA) database (Altenhoff *et al*, 2021) using a Python script written in-house using the PyOMADB client (Kaleb *et al*, 2019) (Kaleb *et al*, 2019) (December 13, 2023, database version July 2023).

### AF-multimer prediction of binary interactions

We used Alphafold-Multimer v2.3.2 (preprint: Evans *et al*, 2021) to generate 3D models of full-length LRRK2 binary interaction complexes for all the interactors shorter than 2000 residues. The databases required to run AlphaFold-Multimer were downloaded on 12 January 2023 and the predictions were run with the flag –max template date = 01-01-1900, to avoid the usage of any available experimental templates. Among the 5 models generated for each LRRK2 binary complex, only the best one was considered for further analysis. The score used to evaluate the models was the default one used by AlphaFold-Multimer (0.2*pTM + 0.8*ipTM).

### Interaction fingerprint

AlphaFold produces probabilities for the C_ß_-C_ß_ distances between residues *i* and *j* to fall into a series of distance bins. For each pair of residues, we calculated the expected value of such distance distribution, yielding a matrix of dimension *l_LRRK2_*x *l_n_*, corresponding to the sequence lengths of LRRK2 and the *n*th interacting partner, respectively. Subsequently, we pooled the distogram on the interaction partner’s side and we assigned to each residue of LRRK2 the minimum expected distance to any residue of the interactor. We refer to this vector of minimum expected distances as the interaction fingerprint of a given interacting protein.

### RMSD calculation

We compared LRRK2 conformations by performing Root Mean Square Deviation (RMSD)-based clustering. To calculate the RMSD we fitted the structures using the C_α_ atoms of the LRRK2 chains from position 983 to the end of the sequence. The N-terminal portion of the protein was excluded because it is very flexible. Calculations were performed using the Superimposer function of the PDB Biopython module (Cock *et al*, 2009) (version 1.83) through customized scripts and pipeline (Matic *et al*, 2023).

### Clustering

The clustering was performed using the Ward method with the *clustermap* function from *seaborn* python library (version 0.11.2). Results were displayed through matplotlib (https://matplotlib.org/) and seaborn (https://seaborn.pydata.org/) libraries using customized python scripts. The number of clusters chosen was one that maximized the mean Silhouette Coefficient of all samples (Rousseeuw, 1987), as computed by the *silhouette_score* function in *scikit-learn* (Pedregosa *et al*, 2011).

## Data availability

The mass spectrometry proteomics data have been deposited to the ProteomeXchange Consortium via the PRIDE (Perez-Riverol *et al*, 2022) partner repository with the dataset identifier PXD048832 and https://doi.org/10.6019/PXD048832 (BioID1, BioID2, miniTurbo), PXD048806 and https://doi.org/10.6019/PXD048806 (LRRK2 proximity-proteomes MLi-2 vs. GZD-524) as well as PXD048808 and https://doi.org/10.6019/PXD048808 (RAB29 co-expression). The BioID interaction network and associated co-evolutionary and structural analysis can be accessed at the following URL: (https://dip.bioinfolab.sns.it/explore/). The code used for the analysis is available at the following links: https://github.com/raimondilab/distogram_analysis, https://github.com/raimondilab/Stable_complex_predictor. A point-to-point protocol for the BioID approach was deposited on Zenodo (DOI: 10.5281/zenodo.17045153).

## Acknowledgments

We are grateful to the staff of the Core Facility for Medical Proteomics at the medical faculty of the University of Tübingen for technical assistance. Furthermore, we gratefully acknowledge computational resources of the Center for High Performance Computing (CHPC) at SNS. CJG received funds from The Michael J. Fox Foundation for Parkinson’s Research (grant ID: MJFF-8068.02, MJFF-8068.04 and MJFF-023920) and iMed – the Helmholtz Initiative on Personalized Medicine.

## Limitations of the study

While the HEK293T system enables large-scale experimental throughput, it may exhibit a cell-type–specific bias, including an overrepresentation of microtubule-binding proteins compared with other cell types. Another limitation is the low endogenous expression of LRRK2 in HEK293T cells. As a result, the study relies on the overexpression of LRRK2 BioID fusion proteins, which may introduce artifacts not present under physiological conditions.

---

**Figure EV1:**
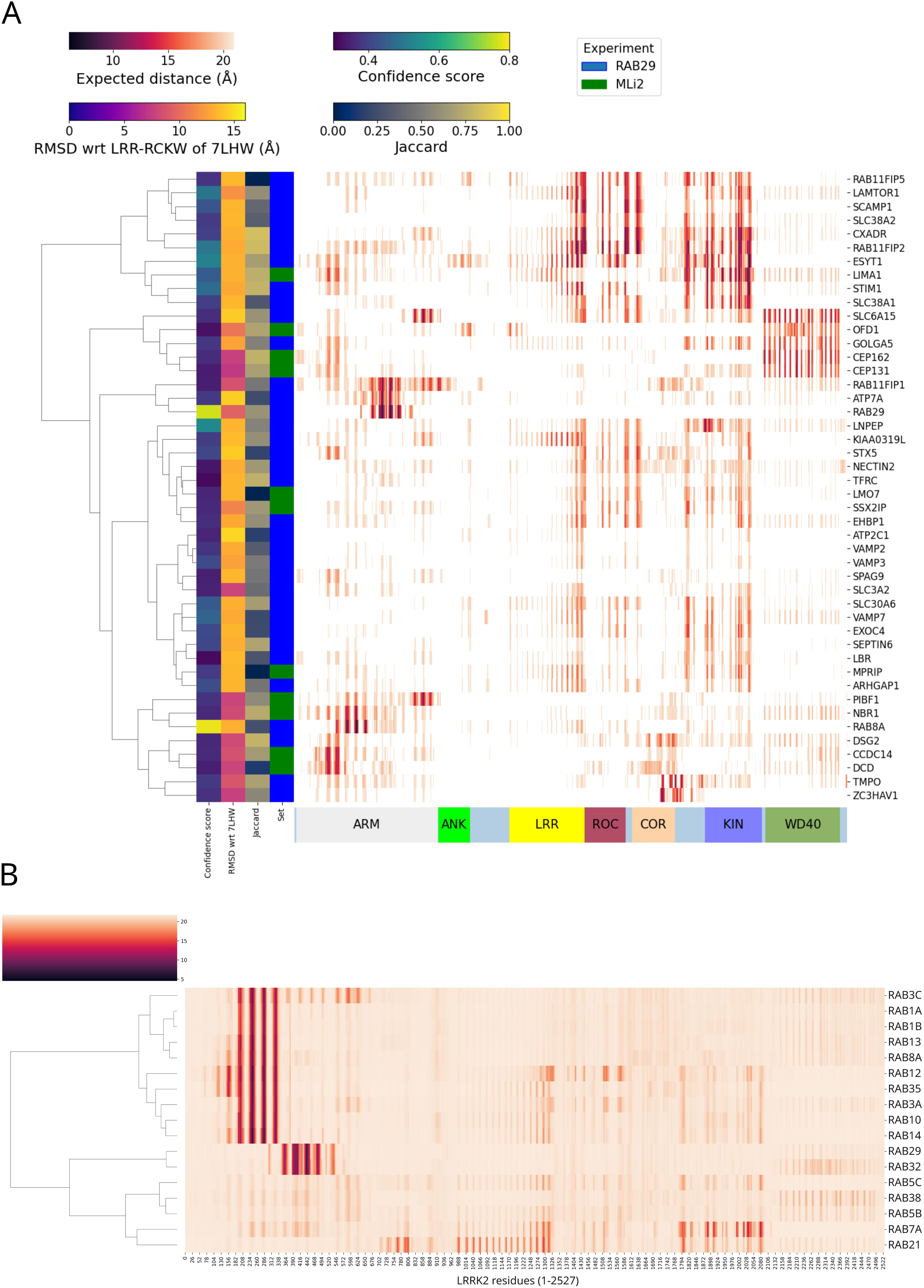
(A) interaction fingerprint clustering of LRRK2’s modulators (MLi-2 and RAB29) interactomes. Cells contain the minimum expected distance obtained from the distograms of any residue of each interactor (row) to every residue in LRRK2 (column). Rows are color annotated based on confidence score, Jaccard, RMSD from reference LRRK2 structure (PDB: 7LHW) and experiment type (either MLi-2 or RAB29 over-expression); (B) interaction fingerprint of LRRK2 and interacting RABs (from IntAct).

## Appendix

**Appendix Figure S1:**
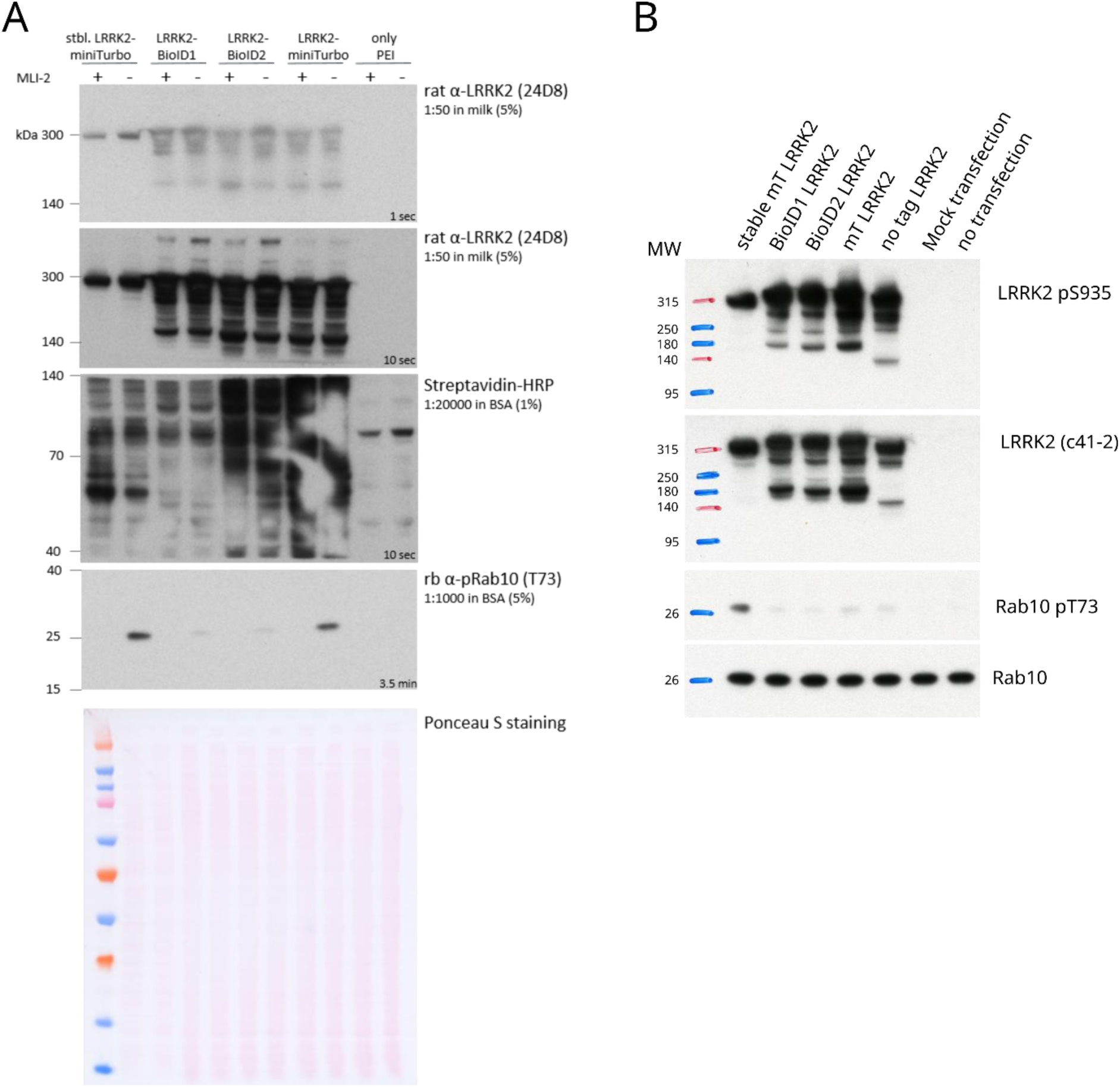
Western blot, pRab10, pS935 and biotinylation – comparison BioID1, BioID2, miniTurbo. To ensure functional BioID-fusion proteins, LRRK2 phosphorylation at S935 and Rab10 phosphorylation was assessed. (A) The different BioID LRRK2 constructs show sufficient biotinylation of proteins as well as MLi-2-responsive Rab10 phosphorylation. (B) All constructs are phosphorylated at S935 (top panel). Replication of LRRK2-mediated Rab10 phosphorylation (last panel) demonstrating functional BioID fusion proteins.

**Appendix Figure S2:**
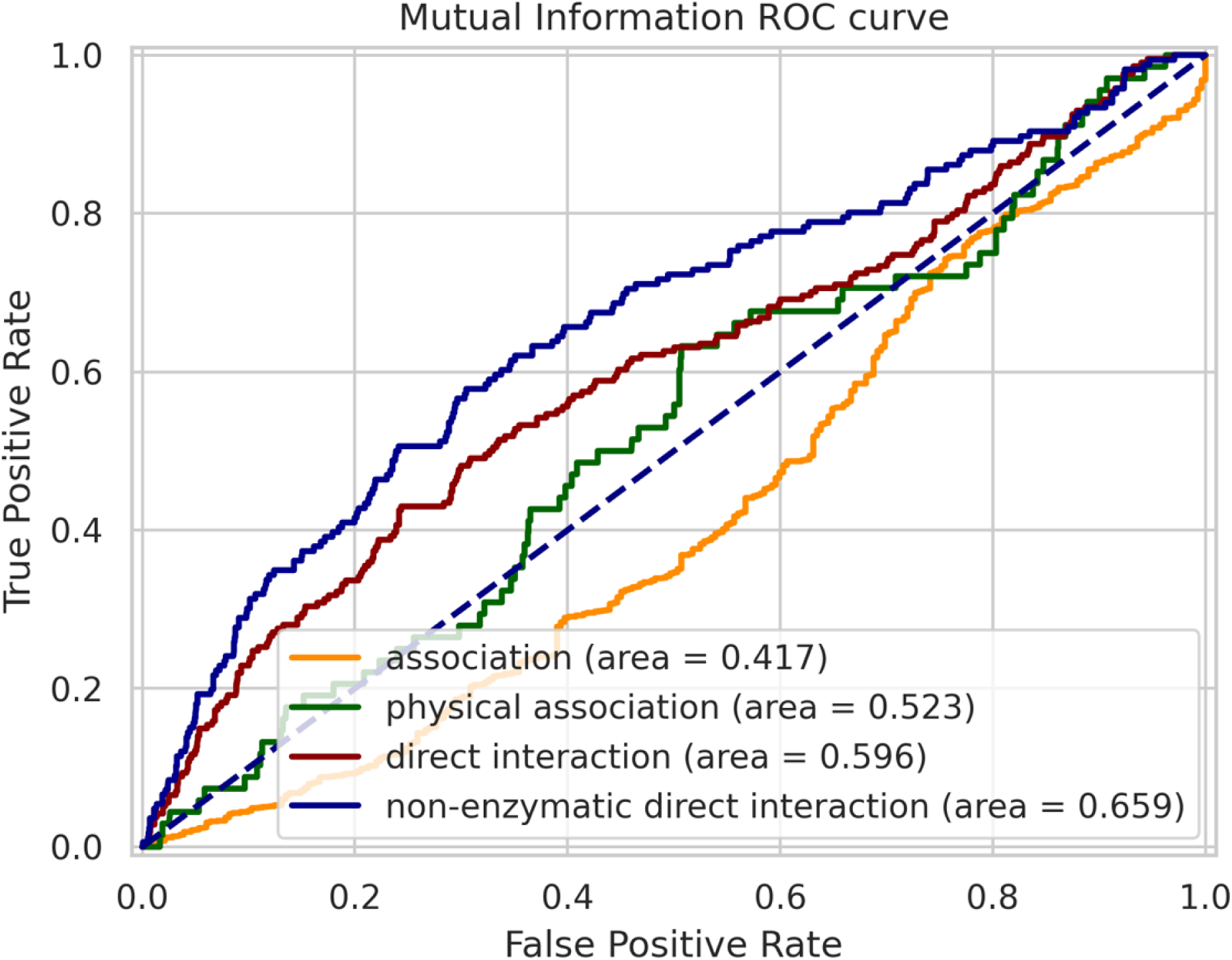
ROC Curve of prediction with Mutual Information (MI) of LRRK2’s IntAct interaction types.

**Appendix Figure S3:**
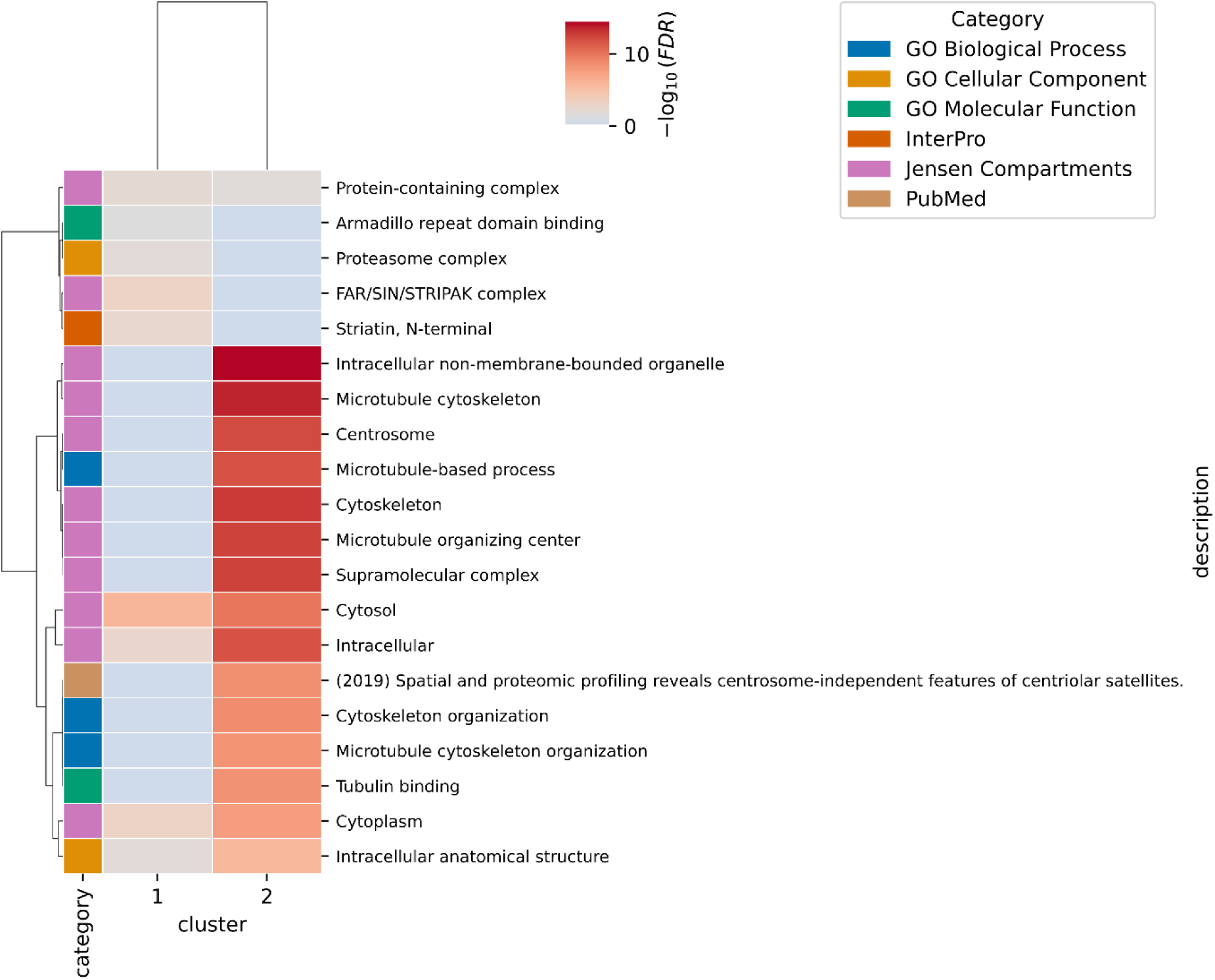
heatmap with cluster specific, significantly enriched processes (FDR < 0.01; StringDB) obtained from co-evolution (Jaccard)-based clustering of the LRRK2 BioID interactome.

**Appendix Figure S4:**
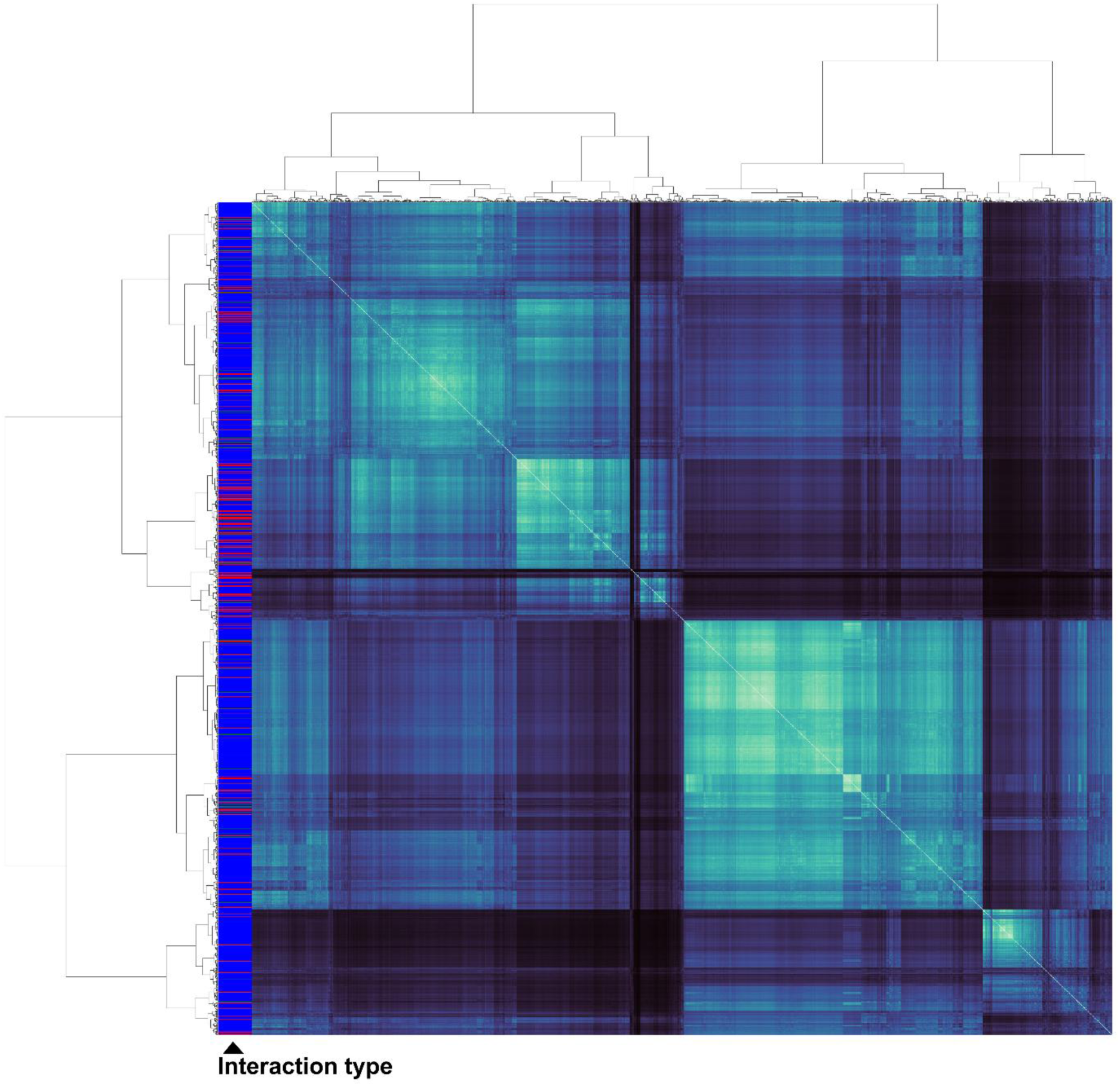
heatmap displaying the coevolution (Jaccard)-based hierarchical clustering of the LRRK2’s IntAct interactome. Colored bars provide IntAct’s interaction type annotation.

**Appendix Figure S5:**
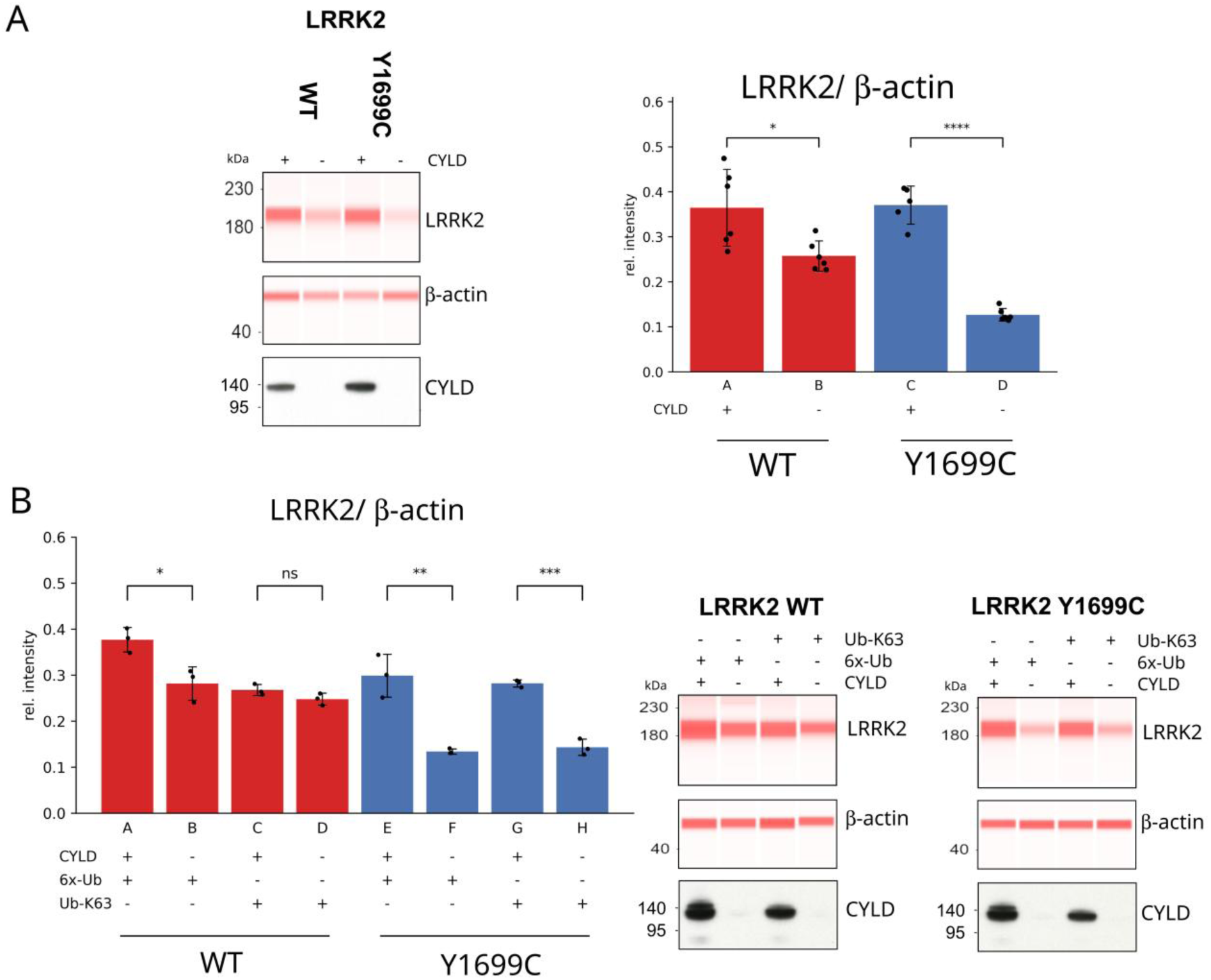
Effect of CYLD co-expression and ubiquitin derivatives on steady-state LRRK2 levels in WT and Y1699C variants (A) Effect of CYLD co-expression on steady-state LRRK2 levels (second independent experiment). LRRK2 WT or Y1699C was co-expressed with CYLD or empty vector control. LRRK2 protein levels were normalized by β-actin. Statistical significance was assessed using unpaired two-tailed t-tests (P-values: A-B: 0.0171; C-D: 3.0E-7; N = 6 biological replicates (for the Y1699C condition, one data point was skipped for technical reasons); error bars = SD) (left panel). Reconstructed Capillary Western signals and confirmation of CYLD expression by Western blot. On the capillary Western blot, LRRK2 migrates at a lower molecular weight (right panel). (B) Effect of ubiquitin conjugation type. LRRK2 WT or Y1699C was co-expressed with CYLD or empty vector control either in the presence of WT ubiquitin or K63-only ubiquitin (Lim *et al*., 2005). Quantification of LRRK2 protein levels from three biological replicates. LRRK2 protein levels were normalized by β-actin. Statistical significance was assessed using unpaired two-tailed t-tests (P-values: A-B: 0.0218; C-D: 0.1289; E-F: 0.0037; G-H: 0.0002; N = 3 biological replicates; error bars = SD) (upper panel). Reconstructed Capillary Western signals and confirmation of CYLD expression by Western blot (lower panel).

**Appendix Figure S6:**
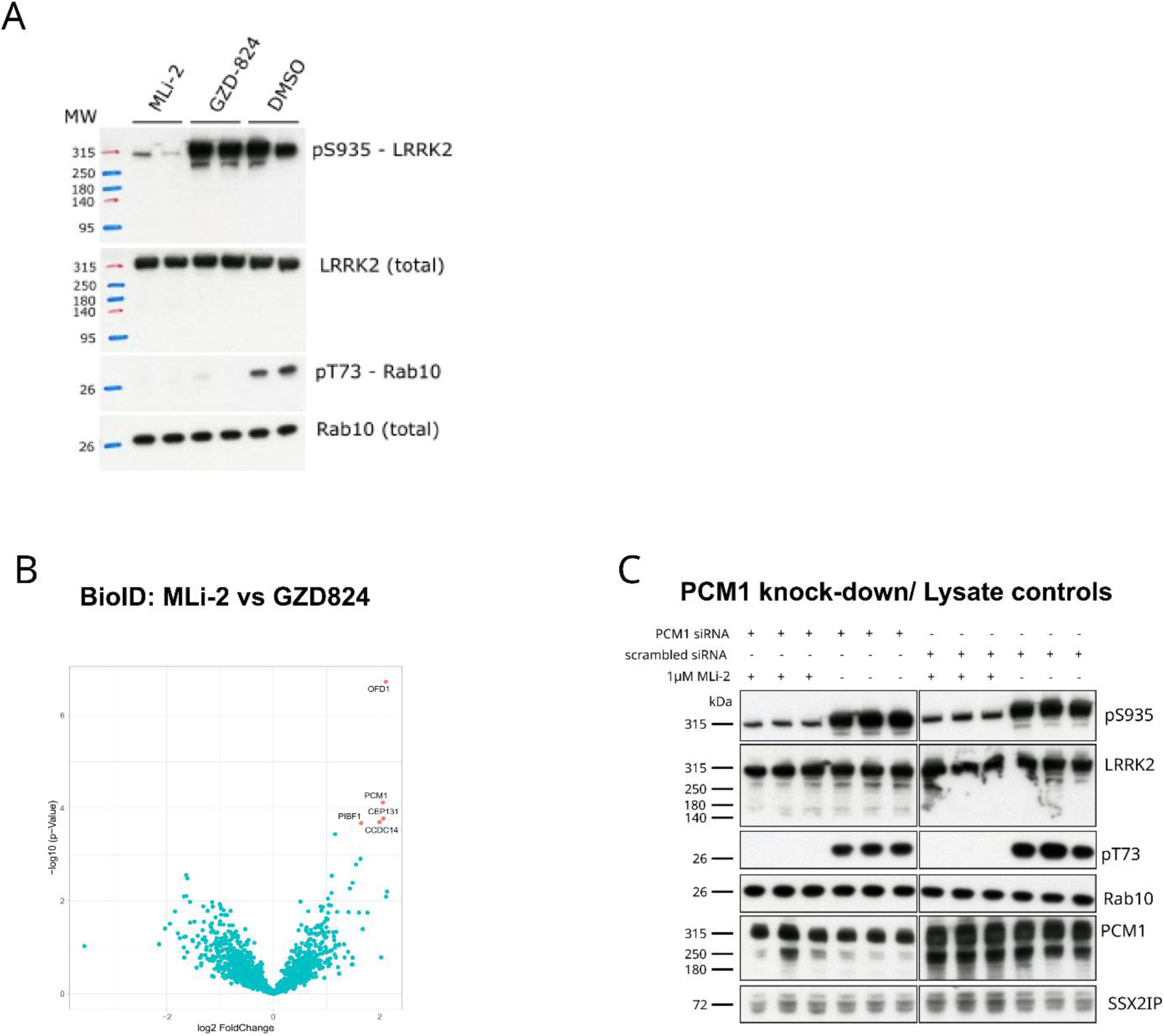
(A) Western blot analysis demonstrating effective LRRK2 inhibition by MLi-2 (type I inhibitor) and GZD-824 (type II inhibitor) in HEK293 cells stably expressing miniTurbo-tagged LRRK2. Both compounds at 1 µM significantly reduce Rab10 phosphorylation. As expected, MLi-2 induces a marked loss of LRRK2 pS935 phosphorylation, a hallmark of type I kinase inhibition. In contrast, GZD-824 does not affect pS935 phosphorylation, consistent with its classification as a type II inhibitor (Tasegian *et al*., 2021). (B) BioID-based proximity labeling comparing the interactome changes upon treatment with MLi-2 versus GZD-824 (moderated permutation-based T-test, FDR = 0.05; S0 = 0.1; N = 7). Volcano plot illustrates differential protein interactions, highlighting the distinct molecular effects of type I and type II LRRK2 inhibition. (C) Validation of PCM1 knock-down by western blot. Efficient depletion of PCM1 is confirmed, with no observed changes in total protein levels of LRRK2, Rab10, or SSX2IP. LRRK2 inhibition by MLi-2 remains effective in the context of PCM1 knock-down.

